# Beyond synthetic lethality: charting the landscape of clinically relevant genetic interactions in cancer

**DOI:** 10.1101/253120

**Authors:** Assaf Magen, Avinash Das, Joo Sang Lee, Mahfuza Sharmin, Alexander Lugo, J. Silvio Gutkind, Alejandro A. Schäffer, Eytan Ruppin, Sridhar Hannenhalli

**Affiliations:** Center for Bioinformatics and Computational Biology, University of Maryland, College Park, Maryland, USA; Cancer Data Science Laboratory, National Cancer Institute, National Institutes of Health, Bethesda, Maryland, USA; Laboratory of Immune Cell Biology, National Cancer Institute, National Institutes of Health, Bethesda, Maryland, USA; Department of Biostatistics and Computational Biology, Harvard School of Public Health, Boston, Massachusetts USA; Massachusetts General Hospital Cancer Center, Harvard Medical School, Boston Massachusetts USA; Department of Genetics, Stanford University, Stanford, California, USA; Moores Cancer Center, University of California San Diego, La Jolla, California, USA

## Abstract

The phenotypic effect of perturbing a gene’s activity depends on the activity level of other genes, reflecting the notion that phenotypes are emergent properties of a network of functionally interacting genes. In the context of cancer, contemporary investigations have primarily focused on just one type of functional genetic interaction (GI) – synthetic lethality (SL). However, there may be additional types of GIs whose systematic identification would enrich the molecular and functional characterization of cancer. Here, we describe a novel data-driven approach called EnGIne, that applied to TCGA data identifies 71,946 GIs spanning 12 distinct types, only a small minority of which are SLs. The detected GIs explain cancer driver genes’ tissue-specificity and differences in patients’ response to drugs, and stratify breast cancer tumors into refined subtypes. These results expand the scope of cancer GIs and lay a conceptual and computational basis for future studies of additional types of GIs and their translational applications. The GI network is accessible online via a web portal [https://amagen.shinyapps.io/cancerapp/].

## Introduction

Cellular functions are mediated by functionally interacting networks of genes. Functional genetic interactions (GIs), whereby the phenotypic effects of a gene’s activity are modified by the activity of another gene, are thus a key to understanding complex diseases, including cancer, which involves an interplay among a myriad of genes (Ashworth et al., 2011; Jerby-Arnon et al., 2014; Kelley and Ideker, 2005; Lu et al., 2013; Wong et al., 2004; Zhong and Sternberg, 2006). GIs are of particular interest in cancer because the dependence of one gene’s phenotypic effect on the activity of another gene provides opportunities for selective killing of cancer cells (Kaelin, 2005) and the interaction partners of drug targets can buffer their effects leading to resistance (Fong et al., 2015; Miyamoto et al., 2015).

In cancer genomics, three types of GIs have been studied so far showing major roles in disease progression and patient survival and suggesting novel therapeutic avenues. The vast majority of GI studies to date have focused on synthetic lethal (SL) gene pairs, describing the relationship between two genes whose individual inactivation results in a viable phenotype while their combined inactivation is lethal to the cell (Ashworth et al., 2011; Miyamoto et al., 2015; Sajesh et al., 2013; Stuhlmiller et al., 2015). They provide selective treatment opportunities by drugs that inhibit an SL partner of a gene that is specifically inactivated or lost in a given tumor, thus selectively killing the tumor cells (Ashworth et al., 2011; Jerby-Arnon et al., 2014; Kroll et al., 1996). Another related class of GIs are Synthetic Dosage Lethal (SDL) interactions, where the underactivity of one gene together with the over-activity of another gene is lethal but not either event alone (Megchelenbrink et al., 2015; Stuhlmiller et al., 2015; Szappanos et al., 2011). SDLs are promising for oncogenes, many of which are difficult to target directly, by targeting their SDL partners (Chang et al., 2013; Luo et al., 2009a; Rathert et al., 2015). A third class of GIs are Synthetic Rescues (SR), where a change in the activity of one gene is lethal to the cell but an alteration in its SR partner ‘rescues’ cell viability. SRs may play a key role in tumor relapse and emergence of resistance to therapy (Brough et al., 2011; Hartwell et al., 1997; McLornan et al., 2014). Indeed, previous investigations have shown that the overall numbers of functionally active SLs and SDLs in a given tumor sample are highly predictive of patient survival (Megchelenbrink et al., 2015). These three interaction types however represent just the ‘tip of the GI iceberg’, as there are many additional types of GI that can be defined at a conceptual level, and whose systematic exploration may have important functional ramifications for cancer therapy.

Here we present a novel data-driven computational pipeline, called “EnGIne” (Encyclopedia of clinically significant GIs in cancer). We applied EnGine to analyze 5,288 TCGA samples (Methods) of 18 different cancer types, identifying clinically significant GIs of 12 distinct types. Using drug response data from TCGA and molecular drug target information, we show that the detected GIs are associated with response to therapy by specific drugs. Their activation patterns can account for the tissue-specificity of known driver genes and stratify breast cancer into clinically relevant subtypes. In sum, EnGIne substantively expands the current knowledge of genetic interactions in cancer, laying a strong conceptual and computational foundation for future studies of additional GI types.

## Results

### Overview of the Encyclopedia of Genetic Interactions (EnGIne) Pipeline

The overall EnGine pipeline is summarized in Fig. 1 and the technical details are provided in the Methods section. Given a large set of tumor transcriptomes (Fig. 1A), we first partition the expression level of each gene into low, medium and high, following our previous approach to identify SL interactions (Lee et al., 2018). Thus, for a pair of genes, there are 9 = 3 × 3 combinations, or bins, of possible co-activity states for the two genes (Fig. 1B). For a given ordered pair of genes, each tumor sample maps to exactly one of the 9 bins. Our goal is to identify GI pairs of the form (x, y, b, ±α) such that for the specific gene pair (x, y), the tumors in which the joint activity of (x, y) maps to bin b have a significant fitness advantage (+) or disadvantage (-) with effect size α, relative to all other tumors whose activity of (x, y) maps to a different bin. The effect size is α estimated by measuring the difference in the survival curves between those patients where the activity of (x, y) is in bin b in their tumors and those where it is not, as depicted in Fig. 1C; note that for most gene pairs, there may not be any bin exhibiting a significant fitness differential. A significant GI pair (x, y, b, ±α) is termed *functionally active* in a particular tumor if the co-activity states of (x, y) in that tumor fall in bin b. We hypothesized that the patients whose tumor has a larger number of functionally active interactions with negative tumor fitness effects will have better prognosis and conversely, the patients whose tumor has a larger number of functionally active interactions with positive tumor fitness effects will have poorer prognosis.

**Figure 1.**
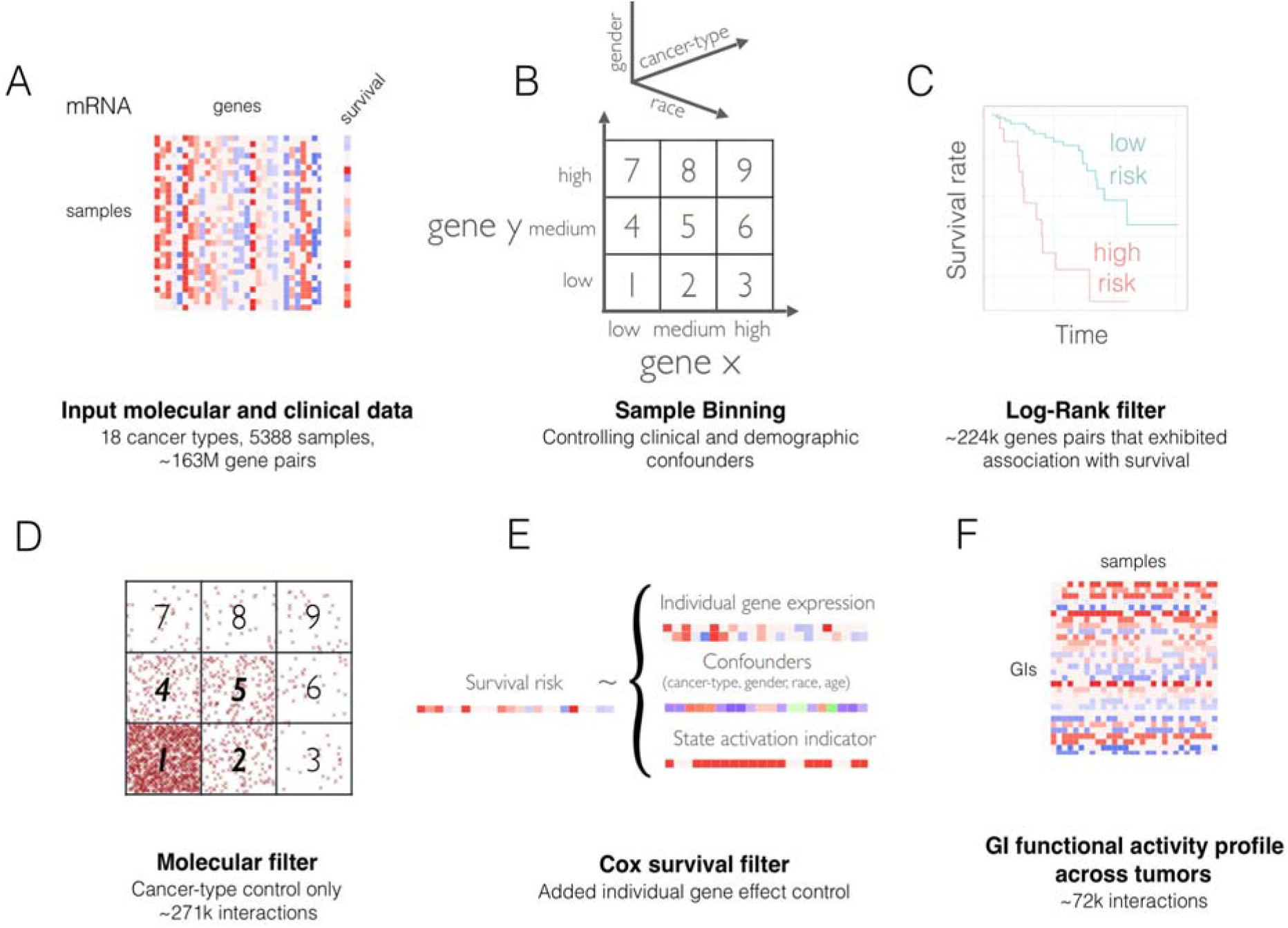
Overview of the EnGIne pipeline. Given a large set of tumor transcriptomes (A), we first partition the expression level of each gene into low, medium, and high activity state, resulting in 9 joint activity state bins for any two genes (B). Each combination of a gene pair and bin *b* induces a bipartition of the set of tumor samples based on whether the co-activity levels of the gene pair in a specific tumor is in bin *b*. The first step of EnGIne screens for the gene pairs that show distinct survival trends in the two sets of tumors in any of the bins, based on log-rank test (C). Next, for a gene pair and a bin identified in (C), we test whether the putative gene interaction in bin *b* has a differential effect on tumor fitness, by testing for depletion/enrichment of samples in the bin *b* relative to expectation based on individual genes (D). Finally, for each retained gene pair, in each of the 9 bins separately, we fit a Cox proportional hazards model to assess whether being in a particular bin is associated with a distinct (positive or negative) pattern of patient survival, followed by correction for multiple hypotheses testing (E). The output of EnGIne is (a) a list of GIs of each of the 12 types studied, and (b) GI profile in each of the individual tumor samples, defined as activity state of each GI in the tumor sample (F).

We analyzed 5,288 TCGA samples for 18 cancer types (see Methods). First, as an initial screening, we performed a Log-Rank survival test (depicted in Fig. 1C) for each gene pair in each of the 9 bins. To make this computationally feasible and to limit the burden of multiple testing correction in the subsequent steps, we used an extremely stringent cutoff for the log rank test leading to the retention of about 1/1,000 gene pairs surveyed (Methods), resulting in 223,946 gene pairs that exhibit a significant association with survival in one of the 9 bins. Second, if a potential GI in bin *b* has a differential effect on tumor fitness, we expect the number of tumors that map to bin *b* to be relatively enriched (for a ‘+’ interaction positively affecting tumor survival), or depleted (for a ‘−’ interaction negatively affecting tumor survival). Thus, we applied an additional filter (Fig. 1D) to retain the GIs exhibiting a consistent patient survival and tumor fitness enrichment or depletion statistic (Methods), yielding 179,444 gene pairs. Third, for each retained gene pair, in each of the 9 bins, we implemented a Cox proportional hazards model, specifically controlling for age, cancer-type, gender, and race, to assess whether a tumor being in a particular bin is associated with patient survival, either positively or negatively (Fig. 1E) (Methods). Finally, we applied an empirical False Discovery Rate (FDR) correction based on the significance of the Cox interaction term of the 179,444 gene pairs relative to those obtained for randomly shuffled gene pairing as the null control. At a False Discovery Rate (FDR) < 1%, this resulted in 71,946 predicted GIs across the 9 bins, of the form (x, y, b, ±α), which form the final set of TCGA inferred GIs (Fig. 1E). Considering the symmetry among bins (bin 2 ~ bin 4 corresponding to low-medium expression interaction; bin 3 ~ bin 7 corresponding to low-high expression interaction; bin 6 ~ bin 8 corresponding to medium-high expression interaction), there are 6 unique types of interaction bins, and considering the two directions of the effect size yields a total of 12 basic types of GIs. We ascertained the robustness of the pipeline to changes in the quantile boundaries for the 3×3 bins and to changes in the log-rank and FDR thresholds (Supplementary note 1).

### The landscape of different GI types

EnGIne identified 71,946 clinically significant GIs of 12 different types, ~0.02% of all the possible candidate gene pairs and GI types tested. Considering the expectation that neighboring genes in the protein interaction network (PIN) are more likely to be involved in a GI (Schaefer et al., 2012), to obtain a smaller but more biologically grounded PIN-supported GI network, we retained only the gene pairs that are separated by two or fewer edges in the human protein interaction network (PIN). This PIN-supported GI network was composed of 1704 GIs involving 1786 genes (Supplementary Table S1) that included 133 known cancer genes (Cosmic dataset) (Futreal et al., 2004) associated with various cancer types (enrichment P-value < 2.5E-22) and 50 breast cancer specific (Intogen dataset) (Gonzalez-Perez et al., 2013) driver genes (enrichment P-value < 7.7E-13).

The distribution of the detected 1704 PIN-supported GIs across the 12 GI types reveals that previously characterized interactions may represent only a small fraction of the overall interaction landscape (Fig. 2A). SL interactions are surprisingly one of the least abundant types of identified GIs, and so are SDLs (1% of all GIs). Remarkably, the positive “anti-symmetric” type of SLs, in which the joint low activity of the two interacting genes is associated with a higher tumor fitness, is 3 times more abundant than SLs. The interaction between the Cosmic Cancer Census genes *GNAQ* and *JAK2* is one example of such a positive interaction in bin 1 (Supplementary Fig. S1). *GNAQ*, encoding Gq, and *JAK2* are both downstream targets in a signaling pathway with several functions pertinent to cancer, including endothelial cell maintenance and vascular remodeling (Kawai et al., 2017). Interestingly, the two most abundant types of pan-cancer GIs correspond to bin 2 and bin 6) where one of the genes has medium level of activity and only the extreme activity of its partner gene reveals a phenotypic effect. For most GI bins, we see a higher proportion of GIs exerting a positive effect on tumor fitness, consistent with the hypothesis that the GIs uncovered during the evolution of cancer are under positive selection. The above distribution trends are quite similar for the full 71,946 GI network (Supplementary Fig. S2A). Additionally, we ascertained that the inferred GIs are not dominated by correlated gene expression patterns (Supplementary note 2).

**Figure 2.**
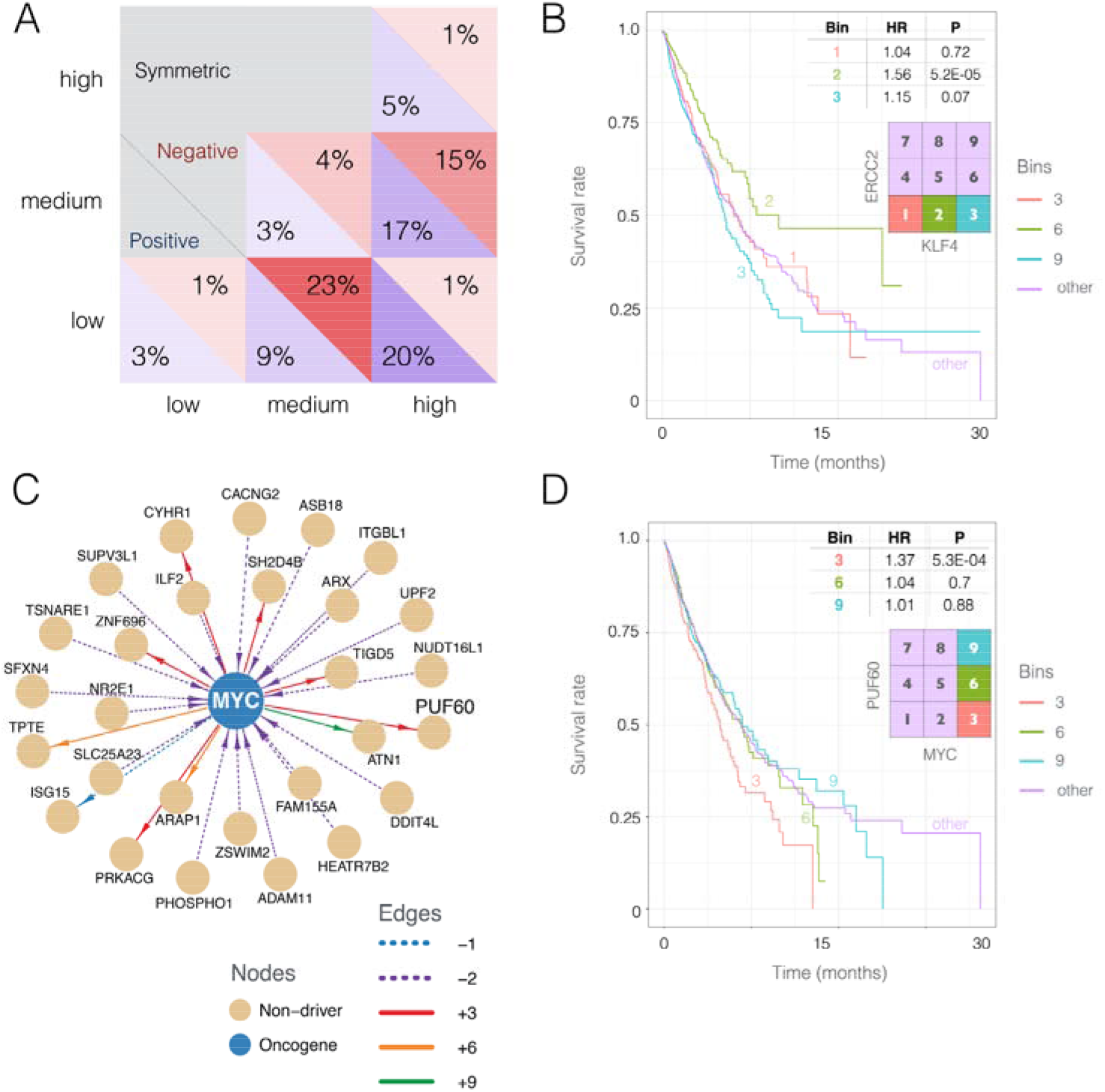
Broad distribution and characteristics of the detected GIs and context-specific effect of cancer driver genes on survival. (A) Distribution of the 1704 significant PIN-supported GIs across 9 joint activity bins. The fractions of GIs in each bin are shown for GIs with positive (blue) and negative (red) effect on tumor fitness. Only the data in the lower triangle of the matrix are shown as the GIs are symmetric relative to the genes in a pair. (B) The Kaplan-Meier (KM) survival curve of GI involving *ERCC2*, a transcription-coupled DNA excision repair gene, known to be a breast cancer tumor suppressor, and *KLF4*, a zinc finger transcription factor known to be oncogenic in breast cancer, reveals increasingly poor survival by over-activation of the oncogene and under-activation of the tumor suppressor (bin 3). Strikingly, the effect of *ERCC2* inactivity on survival is reversed when *KLF4* has medium activity level (bin 2). (C) The predicted GIs involving the oncogene *MYC*. (D) KM survival curve of GI involving *MYC* and its regulator *PUF60*. High expression of *MYC* is associated with poor prognosis specifically at low activation of *PUF60*.

Cancer genes that encode transcription factors, such as *MYC* and *KLF4*, have proven difficult to target directly (Lambert et al., 2018; Li et al., 2018). One important application of EnGIne is to identify candidate interaction partners of the difficult-to-target cancer genes for indirect interventions. To assess this capability, we identified the GI partners of several cancer genes using target-specific FDR (Methods). Fig. 2B shows survival patterns for different activity state combinations of breast tumor suppressor *ERCC2*, a transcription-coupled DNA excision repair gene (Benhamou and Sarasin, 2002; Bernard-Gallon et al., 2008), and a breast cancer oncogene *KLF4*, a zinc finger transcription factor (Akaogi et al., 2009). It reveals two interesting trends: As expected, the over-activation of the oncogene and underactivation of the tumor suppressor (bin 3) results in poorer patient survival than expected from the individual gene effects (bins, 1, 2, 6, and 9). However, surprisingly, the survival curve reveals a reversal of the effect of the tumor suppressor *ERCC2* inactivity on survival when the oncogene *KLF4* has medium activity (bin 2), whose individual activity is associated with better survival; the (*ERCC2, KLF4*) interaction exemplifies the relevance of medium expression bins in this study. This and several other examples of GIs involving a cancer driver gene (Supplementary File 1) demonstrate that the context-specific effects of driver genes may show very different trends than their previously established effects as individual genes. Supplementary Fig. S3 shows the extended GI-network (71,946 GIs prior to PIN filtering) involving the Cosmic and Intogen driver genes. In addition, and consistent with *MYC*’s role as an oncogene, the GIs occurring when *MYC* has low activity mostly have negative effect on tumor fitness. However, when *MYC* is activated, we find that the low expression of *PUF60*, one of the known regulators of *MYC* (Matsushita et al., 2014; Rahmutulla et al., 2013), is associated with higher tumor fitness (type +3 interaction, HR = 1.37, P-value < 5.0E-04) (Fig. 2D). In contrast, we find that high expression of *MYC* does not significantly contribute to poorer prognosis when *PUF60* is expressed at medium or high levels (P-value = 0.9). Thus, this result underscores the importance of molecular context in developing anti-MYC treatments.

We validated EnGIne by comparing its SL predictions to previously reported SLs identified via large *in vitro* screens (Bommi-Reddy et al., 2008; Lord et al., 2010; Luo et al., 2009b; Martin et al., 2009; Steckel et al., 2012; Turner et al., 2008). Each of the three filtering steps of EnGIne (Fig. 1C-E), could discriminate the experimentally determined SLs from the non-SLs, with ROC-AUCs of 0.63 (P-value < 0.0005), 0.62 (P-value < 0.001), and 0.59 (P-value < 0.012), respectively. These results are significant, albeit of modest accuracy (reflecting the widely known discrepancy between in-vitro and in-vivo data (Williams et al., 2000)), support the contribution of each of the individual steps in EnGIne. In addition, we find the PIN-supported GIs to be predictive of patient survival both in cross-validation setting in TCGA (Chang et al., 2013) as well as in an independent breast cancer METABRIC dataset (Curtis et al., 2012) (Fig. S4A, Supplementary note 5). The prediction accuracy, quantified via the concordance index (CI) show that GI based prediction compares favorably with the gene-wise approach (Supplementary results). A bigger improvement is observed in the independent METABRIC dataset (concordance ≈ 0.64), testifying that the GI-based approach is generalizable, while the individual gene-based approach fails to generalize (concordance ≈ 0.51). Supplementary Fig. S4B depicts the survival prediction accuracy of each GI type. Interactions involving both genes in their wild type mid-activity levels (i.e. bin 5) have negligible predictive power on survival, testifying that more extreme levels of expression of at least one of the two genes tend to be involved in functional GIs affecting survival.

### Differential activity of drug target GIs between responders and non-responders

Aiming to test the ability of EnGIne inferred GIs to predict drug response, we applied EnGIne to identify GIs based only on TCGA samples that do not have drug response information, and tested the predicted GIs’ ability to discriminate responders from non-responders in the ‘unseen’ TCGA samples where the drug response information is available (Methods). Notably, because the considered drugs are inhibitory, it suffices to focus on GI bins 1, 2, and 3, where one of the genes (the drug’s target) has low activity. For a given drug and cancer type having data on responders and non-responders, we analyzed the GIs involving each of the drug targets (identified via target-specific FDR; Methods). We then tested whether the frequencies of GI activation in responders and the non-responders are significantly different using a Fisher exact test (Fisher, 1922) (Methods). For positive GIs, we expect a lower GI activation frequency among responders and the opposite for negative GIs (e.g., as in the case of SL-type GIs). However, owing to very small and unbalanced numbers of responders and non-responders (5 to 35 samples per response group per drug), the Fisher test is underpowered, and we therefore, compared the Fisher test p-values of the GIs (equivalently, ratio of GI frequency in responders and non-responders), segregated over all GIs of a specific type, with those obtained using randomly shuffled drug-response labels, using paired Wilcoxon tests (Wilcoxon, 1945) (Methods), performed separately for each drug-cancer type pair.

We considered the 12 drug-cancer type pairs that have RECIST (Response Evaluation Criteria In Solid Tumors) (Eisenhauer et al., 2009) drug response following treatment for at least 10 patients (at least 5 responders and 5 non-responders) in TCGA. Each of the 6 basic GI types was tested for the 12 drug-cancer type pair. Overall, in 18 of the 72 tests (5 fold enrichment for P ≤ 0.05) of drug-cancer type combinations, GIs of a particular type exhibit statistically significant differential activation frequencies between responders and non-responders consistent with the expected effects of the GIs (Fig. 3A). Reassuringly, several of those significant drug-target GI’s are in bin 1, which contains the SLs, consistent with previous reports showing the role of SLs in mediating drug response (Jerby et al.). Among the drugs, Gemcitabine, Lomustine, and Paclitaxel exhibit differential GI activation for most GI types (aggregate P-values ranging from 4E-16 to 2E-11). We also explored the most differentially activated individual GIs. Imposed an empirical FDR threshold of 0.01 on the Fisher test P-value yielded 521 GIs for the 12 drug-cancer type combinations (Methods, Supplementary Table S2). Individual genes comprising the 521 GIs are closer to each other in the PPI network relatively to shuffled pairs (Wilcoxon P < 0.001, Methods) and have a significantly increased number of direct PPI interactions between them (Fisher P < 0.02, Methods).

**Figure 3.**
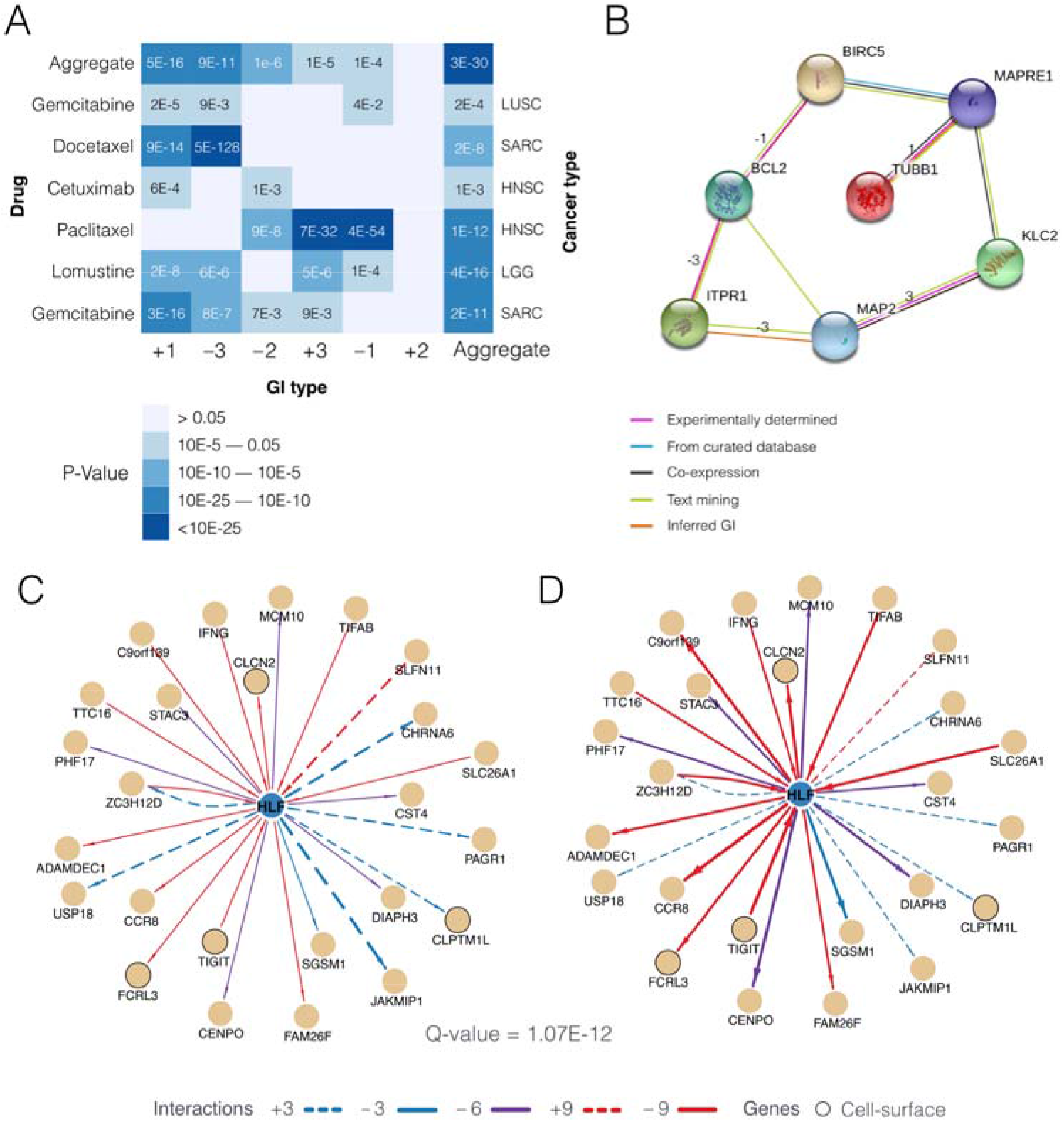
(A-B) differential GI activation between drug response groups. (A) For each drug (left row labels) and each cancer type (right row label) combination, and for each GI type (columns), the heat plot shows the significance of differential activation of GIs in responders and non-responders consistent with expectation. The last column shows the significance when all GI types are aggregated. (B) The network shows the inferred functional interactions (based on STRING (Szklarczyk et al., 2015)) among the genes interacting with *Paclitaxel* targets, as well as inferred GI types. **(C-D) Differential GI activation between tissues in gene-specific GI-network.** For the HLF-specific GI network, the figure shows the activity states of GIs in breast and lung cancers (C - foreground tissues) and in other cancer types (D - background tissues). The edge weight (thickness) represents the fraction of samples in which the GI was functionally active. Several GIs can be seen as differentially active in the two sets of cancer types. The figure also depicts cell surface proteins among the HLF’s GI partners. The GI network-based sample-specific risk score is significantly higher (q-value < 1.07E-12) in breast and lung cancer relative to other cancer types, potentially mediated by a selective activation of positive GIs in the foreground tissues and negative GIs in the background tissues.

As an illustrative test case, we explored the GIs associated with the response to Paclitaxel, in TCGA Head and Neck Squamous Cell Carcinoma (HNSC) cohort. Paclitaxel inhibits the proteins encoded by *BCL2, TUBB1*, and *MAP* based on DrugBank (Law et al., 2014). We identified a GI involving the inactivation of *BCL2* (known to suppress apoptosis, indirectly inhibited through phosphorylation (Ruvolo et al., 2001)) and the over-activation of *ITPR1* (Inositol 1,4,5-trisphosphate receptor type 1, also known as IP3 receptor type 1), negatively affecting tumor fitness (GI type −3). Interestingly, our analysis shows that this GI is functionally active among the responders at a significantly higher ratio than among non-responders (odds-ratio ≈ 11.1). The interaction between ITPR1 and BCL2 is well characterized (Chen et al., 2004; Oakes et al., 2005; Rong et al., 2009); one of these studies suggests that BCL2 also interacts with the two other human paralogs ITPR2 and ITPR3, but these interactions are not represented in the PIN used in this study and were therefore not detected. BCL2 exerts its oncogenic effect by inhibiting ITPR3-mediated channel opening and Ca^2+^ release from the endoplasmic reticulum, and thus preventing cancer cell apoptosis. Our analysis strongly suggests that BCL2 inhibition by Paclitaxel is especially effective when the ITPR1 expression is abundant, enabling effective Ca^2+^ release. Additional Paclitaxel targets *TUBB1* and *MAP2* are also linked with ITPR1/BCL2 through GIs with literature evidence for experimentally validated or putative interactions (by STRING DB (Szklarczyk et al., 2015)) (Fig. 3B), suggesting promising avenues for additional studies.

### GIs can explain why some cancer driver genes are implicated in some cancer types and not in others

Many of the known cancer driver genes affect tumor initiation and development in a tissue-specific manner, despite the cancer gene being expressed in other tissues as well. Next, we explored whether the GIs can explain the tissue-specificity of cancer genes. Toward this, we identified 15 oncogenes and 20 tumor suppressors whose effects are likely to be restricted to specific cancer types, based on preferentially high mutation rates in those cancer types, including breast, bladder, and gastric cancer (Methods, Supplementary Table S3). For each cancer driver, we assigned a risk score to each patient by aggregating functionally active GIs involving the driver gene defined using target-specific FDR (Methods); for oncogenes, only the bins with high oncogene activity and for tumor suppressors, only the bins with low tumor suppressor activity were considered (Methods). We hypothesized that for a cancer gene, the risk score will be greater in tissues where the cancer gene is implicated relative to other tissues. Indeed, for 15 out of 35 (~43%; 5 oncogenes and 10 tumor suppressors) driver genes, the observations are consistent with our hypothesis (Wilcoxon rank-sum test, FDR < 0.1, Supplementary Table S3).

For instance, HLF, a bZIP transcription factor, has been linked to lung and breast cancer based on its significantly greater missense mutation frequency in those cancer types (Gonzalez-Perez et al., 2013). We observed a significant difference (FDR < 1.07E-12) in GI activation risk score for breast and lung cancer relative to the other tissues. Specifically, we found that positive GIs are preferentially activated in these two tissues while negative GIs are preferentially activated in the other tissues, consistent with the increased tumor fitness in these two foreground tissues (Fig. 3C,D). Overall, these results suggest that cancer type-specific effects of many driver genes may be explained by their tissue-specific GI network activity.

### Stratifying breast cancer tumors into distinct sub-types based on their GI profiles

Next, we investigated whether functionally active pan-cancer GIs in a tumor may provide an alternative methodology to tumor stratification into sub-types. We focus on breast cancer because it has a large number of samples in TCGA and because a second independent dataset, METABRIC, is publicly available. We represent each sample by the functional activity (a binary indicator) of each PIN-supported GI detected in TCGA, rather than generating BRCA-specific network, thus avoiding potential circularity of inference and prediction within samples sharing similar characteristics. Based on this 1704dimensional binary vector representation of each tumor sample we clustered the 1981 breast cancer samples in the independent METABRIC dataset (Curtis et al., 2012) using a conventional Non-Negative Matrix Factorization (NMF) (Methods). Optimal clustering was achieved (maximum value of the Dunn index, Methods) for 10 clusters (Supplementary Fig. S7A). Upon closer inspection of the distributions of known breast cancer subtypes in these clusters (Supplementary Fig. S7B) we merged two of the clusters, thus yielding 9 clusters for further analyses.

Kaplan-Meier curves (Fig. 4A) and statistical analysis show that the 9 clusters have distinct survival characteristics with an overall mean hazard ratio (HR) difference of 1.94 (P-value below the lowest reportable threshold and shown as 0). The distinct survival characteristics are consistent with analysis performed using the full 71,946 GI network (Supplementary Fig. S7C). Supplementary Fig. S8A shows the survival characteristics obtained for the previously published clustering of the METABRIC samples (Curtis et al., 2012). As evident, both approaches obtain similar survival separation levels, but exhibit differences in their histopathological composition. Currently, breast cancer has 5 well-established clinically distinct subtypes based on the tumors’ histopathological attributes. Fig. 4B shows the fractions of each known subtypes among the 9 GI-based clusters. Several clusters are highly associated with specific subtypes such as Basal [triple-negative] (cluster 5), Luminal A (clusters 3,4) etc. Others show association with several subtypes, e.g., Luminal A and B both have high fractions in cluster 8. Interestingly, the basal subtype, which are largely triple-negative and have poor prognosis, correspond to a distinct cluster in our analysis (cluster 5), consistent with their distinct clinical status. In the original METABRIC publication (Curtis et al., 2012), 50% of the samples were left unassigned to any of their 10 clusters, while our GI-based clustering covers all samples.

**Figure 4.**
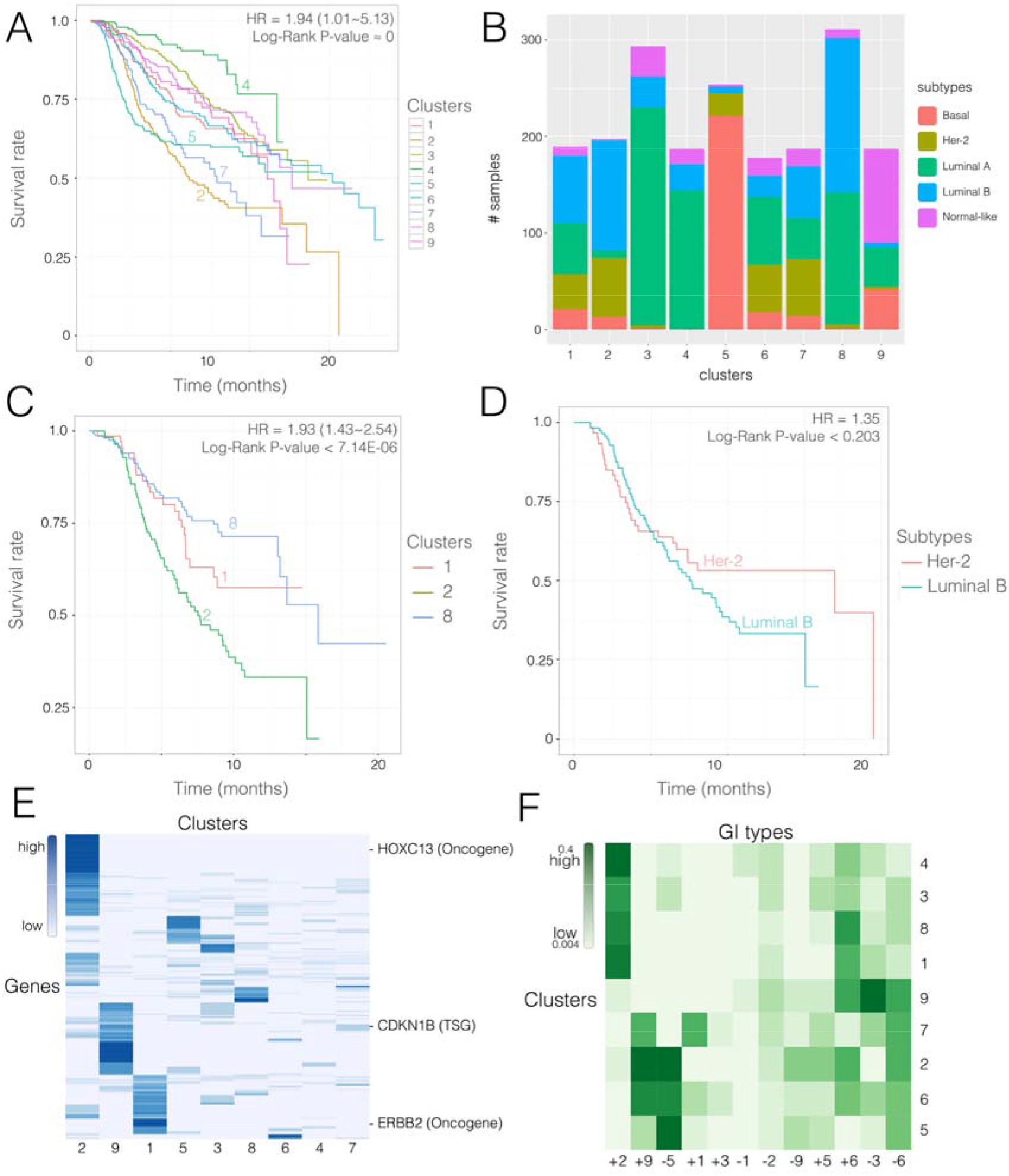
**(A) Mean survival curves of the individuals in the 9 inferred GI-based breast cancer subtypes. (B) Cluster subtype composition based on PAM50 breast cancer sub typing** (Bernard et al., 2009). **(C-D) Survival trends of tumors of known histopathological cancer subtypes within and across GI-based clusters**. (C) Luminal B samples that are split across GI-based clusters 1, 2 and 8 show significant survival differences. (D) Her2 and Luminal-B type tumors that are included within the GI-based cluster 2 exhibit similar survival trends. **(E) Mutational profile of GI-based breast cancer subtypes**. Mutation profiles of 196 genes (rows) across the 9 GI-based clusters (columns). For each gene and each cluster, the figure depicts the fraction of samples in the cluster in which the gene is mutated. **(F) GI types composition of the GI-based breast cancer subtypes in the METABRIC dataset**. In clustering the samples based on GI profile, each GI is probabilistically assigned to a single cluster, based on which, the composition of GIs assigned to each cluster is obtained. The x-axes represent the 12 GI types (6 activity bins and 2 directional effects on survival), and the y-axes represent the clusters. The colors represent the fraction of cluster-assigned GIs of each GI type.

We find the GI based approach to provide improved survival predictive value over the classical histopathological ones (Fig. 4C,D). There are two situations in which the GI-based clustering leads to a different classification of patients for survival analysis: (1) cases where known histopathological breast cancer subtypes are split across multiple GI-based clusters (e.g. Luminal B across clusters 1, 2 and 8), and conversely (2), cases where one GI-based cluster harbors multiple known histopathological subtypes (e.g. cluster 2 contains Her-2 and Luminal B subtypes). In the former case, we find that the 1989 Luminal B tumors that are split across different GI-based clusters exhibit statistically significant (P < 7.14E-06) distinct survival trends (Fig. 4C), supporting the GI approach in separating the Luminal B tumors. Likewise, in the latter case, we find that the survival trends of Her-2 and Luminal B histopathological subtype samples that are assigned to the same GI-based cluster 8 do not show a significant difference (P < 0.203) in their survival trends (Fig. 4D), suggesting that the GI-based stratification may in some instances provide better survival prognosis relative to histopathology-based stratification. We systematically identified 6 additional instances of the above two scenarios where (1) a known tumor histopathological subtype was split across multiple GI-based clusters or (2) multiple known histopathological subtypes were assigned to the same GI-based cluster (and each cluster has at least 30 samples). In each instance of the first kind we tested for statistically significant differences in survival and in each instance of the second kind we tested for lack thereof. As shown in Supplementary Fig. S9, in 5 out of 6 instances we found that the GI-based clusters provided a more accurate survival prognosis. In contrast, we identified 4 cases of the second kind in the original METABRIC clusters (Curtis et al., 2012) and found that none of their survival trends were significantly different (Supplementary Fig. S10). Thus, these results demonstrate that the GI approach performs better than clustering based on histopathological subtypes or METABRIC clustering based on gene expression profiles, in terms of survival prognosis.

To explore potential mutational basis of the GI-based clusters in another way, we assessed whether the samples in GI-based clusters harbor distinct mutations patterns. We identified 196 genes (Supplementary Table S4) with significantly greater mutation frequency in one or more of the clusters, relative to their overall mutation frequency in breast cancer (Methods). Fig. 4E shows the mutational frequency profiles of these genes across the 9 clusters. Overall, the differentially mutated genes across the GI based clusters include 10 cancer drivers: *CDK12, CDKN1B, DNAJB1, ERBB2, EXT2, FCGR2B, FNBP1, HOXC13, PDGFRB*, and *SEC24D*. A more detailed discussion of the potential biological significance of some of these mutations is provided in the Supplementary Results. We note that no mutation data was used in the GIs inference via EnGIne.

Finally, we quantified the fraction of each of the 12 types of functionally active GIs among the samples in each cluster (see Methods). Fig. 4F shows the active GI profiles of each cluster, and reveals two broad subgroups, one including clusters 4, 3, 8, and 1 and another including clusters 2, 5, 6, and 7. Interestingly, the two subgroups clearly segregate in terms of their survival, testifying that the classification into GI types captures a simplified yet robust characterization of the clinical prognosis. The first broad subgroup of tumors (clusters 4, 3, 8 and 1) are characterized by high fractions of type +2 and +6 GIs, both of which involve a medium expression and low expression bin. Therefore, this analysis demonstrates the relevance of considering medium expression states in molecular stratification. Supplementary Fig. S11 compares the GI-profiles of clusters revealed in the TCGA and the METABRIC breast cancer data and shows a high degree of consistency. A global comparison of the GI profiles of the 9 clusters in the two datasets shows a Spearman correlation of 0.67 (P-value < 2.4E-14) between the GI types composition of these clusters, implying that GI-profiles are a robust characteristic of breast cancer tumors across different tumor collections. Thus, the GI-based clustering demonstrates a proof of principle for improved stratification of breast cancer tumors into classes with distinct survival prognosis and mutational profiles.

## Discussion

Analyzing molecular and clinical data across thousands of tumors of dozens of types, here, for the first time, we comprehensively map the landscape of 12 basic GI types in cancer. Our work extends previous investigations of gene interactions in cancer, which have been almost entirely focused on synthetic lethality (SLs, corresponding to positive effect on survival in bin 1), with a few studies of synthetic dosage lethality (SDL: corresponding to bins 3 and 7), to a total of 12 types of interactions. The identified GIs are predictive of patient survival and drug response, explain tissue-specificity of cancer driver genes, and reveal novel functionally and clinically relevant breast cancer subtypes. The set of functionally active GIs thus provides a complementary molecular characterization of tumors to those obtained by histology and contemporary individual gene-centric transcriptomic and sequence-based profiles. A better concordance of the GI-based breast cancer subtypes with survival trends may be partly because GIs were inferred based on their impact on survival. However, interestingly, the detected subtypes are additionally marked by distinct mutational profiles, which were not utilized in inferring the GIs. Overall, these results underscore the importance of molecular context represented by functionally active GIs.

Multiple factors are worth considering when deciding on the strategy to bin tumors into bins based on gene activity levels. For instance, a previous study of synthetic lethality in glioblastoma (Szczurek et al., 2013) defined the high (low) state as mRNA expression higher than the 80^th^ quantile (lower than the 20^th^ quantile, respectively). We chose to partition gene activity into 3 quantiles of low, medium, and high activity levels within cancer samples. We have shown the robustness of the detected GIs (Supplementary note 1). Besides its simplicity and being non-parametric, our strategy naturally allows us to search for cases where even the normal (or medium) levels of expression of a gene may be associated with fitness effects, depending on the states of other genes (e.g, bins 4 and 7).

Identifying pairwise gene interaction is only a first step toward capturing the true complexity of cellular networks. Future work can go beyond the 12 basic GI types studied here to investigate more complex GI types that involve different compositions of these basic types; for instance, a given interacting gene pair can confer tumor fitness benefit in multiple co-activity bins and reduce tumor fitness in others. Thus, while the results presented here go markedly beyond previous definitions and analyses of GIs, they only begin to explore the full scope and clinical potential of GI-based analyses of cancer, awaiting future investigations.

## Methods

### Cancer datasets

We downloaded The Cancer Genome Atlas (TCGA) (Chang et al., 2013) molecular profiles and clinical covariates via the Broad Firehose (https://gdac.broadinstitute.org/, downloaded on Jan 28, 2016). This covers RSEM-normalized RNAseq data, mutation, and clinical information such as age, sex, race, tumor types, and overall survival of the 8,749 patients (data quality testing is described in Supplementary note 3). Drug response information was downloaded from TCGA data portal available in the form of RECIST criteria (Eisenhauer et al., 2009) and mapped using DrugBank (Law et al., 2014) database V4.0. For the drug response analysis, to consider only gene inactivation mechanism, we excluded those drugs whose DrugBank mechanism of action label is either potentiator, inducer, positive allosteric modulator, intercalation, stimulator, positive modulator, activator, partial agonist, or agonist.

We performed gene expression binning specifically for each combination of cancer type, race and gender. Furthermore, we control for various clinical and demographic group specific effects. Because using small groups of samples may result in insufficiently robust models, we filtered rare combinations of clinical and demographic groups, resulting in 5288 mRNA samples derived from patients spanning 18 cancer types, 3 races and 2 genders. The data were not stratified by stage and grade for three reasons: 1) the grade is missing for most cancer types 2) the stage/grade system varies across tumor types 3) further stratification would result in further loss of data due to small group sizes. We applied quantile normalization within each sample of the expression data. The METABRIC breast cancer dataset (Curtis et al., 2012) (as described in reference (Jerby-Arnon et al., 2014)) consists of 1989 microarray samples and was used for independent validation. Similarly, quantile normalization was applied in each sample.

### Protein interaction Network (PIN)

The PIN was obtained from a previously published resource called HIPPIE (version 2.0, http://cbdm-1.zdv.uni-mainz.de/~mschaefer/hippie/), which aggregates physical protein interaction data from 10 source databases and 11 studies (Schaefer et al., 2012). This network consists of 15,673 human proteins and 203,159 interacting pairs.

### Identification of gene interactions (GI) associated with cancer patient survival

As shown in Fig. 1, we divided the range of each gene’s expression across tumor samples into 3 equal-sized bins that correspond to the 3 activity states: low, medium and high expression. Given a gene pair, each tumor sample is thus mapped into one of the 9 joint activity states of the two genes. The choice of dividing a gene’s activity into three classes, while somewhat arbitrary, was made in consideration of interpretability of functional states and robustness of inference. However, to account for differences in expression distributions across clinical and demographic confounders, we apply a subpopulation specific binning approach. We considered the following categorical confounders: cancer type, race, and gender. We considered the combination of confounder states for which there were at least 100 tumor samples (Supplementary Table S5). Our binning is thus not confounded by various clinical and demographic variables.

The GI pipeline consists of three steps that successively refine the predictions to arrive at high-confidence set of predicted GIs. As such, the number of all pair-wise combinations of genes is excessively large to apply a comprehensive Cox regression model. For the principal analysis shown in the manuscript, specific parameter thresholds were chosen to make the subsequent analysis tractable, but users of the GI pipeline may choose other thresholds. To inform such decisions, we did a robustness analysis of the parameter settings with a smaller input set of gene pairs (Supplementary note 1).

#### Step 1: Log-Rank

In the first step, for each of the ~163 million gene pairs, say (x,y), we compute the Log-rank statistics (Harrington and Fleming, 1982) estimating the survival difference between the samples that map to one of the 9 activity bins and the other 8 bins. We implemented the log-rank test in C++ for computational speed. To control for gene-wise effect, we compare the Log-rank statistics of the gene pair (x,y) (in a bin) with those for (x,U) and separately with those for (V,y), where U and V represent all other genes. For a candidate gene pair (x,y), we consider Log-rank(X,Y) to be significant if it is among the top 0.1% relative to all (x,U) and top 0.1% among all (V,y) gene pairs. This threshold of 0.1% (1/1000) can be controlled by the user. We retain a gene pair if it is deemed significant in any of the 9 bins. This procedure retained 223,946 gene pairs of the total of ~163M.

#### Step 2: Molecular enrichment and depletion

For a gene interaction having positive (respectively, negative) effect on survival, we expect the tumor having that interaction to be under negative (respectively, positive) selection, and therefore we expect the fraction of such tumors (i.e., those mapped to the corresponding activity bin relative to the interacting gene pair to be depleted (respectively, enriched). We only retained the potentially interacting gene pairs for which the fraction of samples in a particular bin, suggested by the log-rank test, were lower (bottom 45 percent) or respectively, greater (top 45 percent among all gene pairs) than expectation, reducing the number of GIs to 271,096 across 179,444 gene pairs. Recall that a gene pair can participate in multiple GIs corresponding to multiple bins and effect on survival. The threshold of 45% can be controlled by the user. Our choice of threshold ascertained that molecular enrichment/depletion is consistent with log-rank test without being overly punitive.

#### Step 3: Cox proportional hazard test

The Cox proportional hazards model is the most widely accepted approach for modeling survival while accounting for censored data as well as confounding factors. For each gene pair passing the filter at step 2, we modeled its effect on survival in each of the 9 bins using the interaction status *λ* (active if the sample mapped to the bin and inactive otherwise), along with the confounders. Specifically, we introduced the expression levels of the two individual genes *σ*_1_, *σ*_2_ to model each gene’s independent effect on survival, and additionally, clinical and demographic confounders, namely, cancer type, race, gender, and age. The model is stratified based on the discrete confounders, to account for differences in the baseline hazard (risk) characteristics. We did not control for tumor stage and grade as these classifications reflect the very same tumor characteristics our model aims to capture, and such control would prevent us from learning an important element of the disease. Control for genomic-stability and tumor purity as a potential confounder is described in Supplementary note 4.

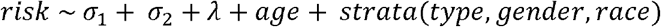

Cox modeling provides a p-value representing the significance of the effect of joint gene pair activity on survival. To obtain a null distribution for the P-values, we repeated this process for corresponding list of randomly shuffled pairs (only among the pairs qualifying step 2 above). We retained ~71K gene pairs in the above the most significant 99^th^ quantile of the null p-values distribution as an empirical FDR control.

#### Optional Step 4: Filtering by protein interactions

To gain additional confidence in the predicted GIs, given the greater tendency (and expectation) for neighbors in the Protein Interaction Network (PIN) to exhibit functional interactions (see Results), we further refined the GI set by retaining the pairs that are found within distance of 2 in the HIPPIE PIN. Overall, we obtain a set of 1704 GIs that exhibit molecular and clinical evidence in cancer as well as evidence from the PIN network.

The pipeline is implemented in R and C++ in a distributed computing environment using SLURM (Yoo et al., 2003) to run many jobs in parallel on a computer cluster.

### Survival Risk Score Computation

We applied a semi-supervised approach to assign a risk score to each patient according to the functionally active GIs in the sample. Consider a GI involving genes x and y conferring a positive effect on the tumor fitness in a particular bin B (Fig. 1). If in a sample, genes x and y fall in bin B, then the GI is said to be ‘functionally active’ in the sample, and a score of +1 is contributed to the overall tumor ‘fitness’. Likewise, if the GI has negative effect on the tumor fitness, then a score of −1 is contributed. The overall risk score given a set of GIs is the sum of the individual GI +1 or −1 scores.

### Patient Survival Risk Prediction

For each sample, we computed the overall score conferred by functionally active GIs (either in a bin-specific and effect direction-specific fashion, or overall) in the sample. The higher the tumor ‘fitness’ score the lower the survival potential. However, to make our approach comparable to gene-wise approaches (Yuan et al., 2014), we assigned each gene the sum of the contributions by all active GIs involving that gene, with multiplicity for gene pairs involved in multiple GIs. The estimated gene-wise GI score is used as a predictor variable in a Cox model along with the confounding factors discussed above to predict patient’s survival.

For cross validation, this model was trained on the same data used for the GIs training and then validated on its cross-validation counterpart. For independent validation, the model was trained on the full TCGA dataset, and tested on the independent METABRIC breast cancer data with 1989 samples (Curtis et al., 2012). The prediction accuracy is estimated in terms of the C-index (Harrell et al., 2005). Several previous publications have assessed survival risk prediction accuracy based on dichotomized analysis where samples are separated into distinct low- and high-survival groups and their survival curves then compared (Harrington and Fleming, 1982), which is prone to overestimating prediction accuracy. For comparison, we also performed accuracy estimation following the dichotomized comparison of survival risks between the extreme cases of predicted survival risk groups, for variable thresholds to define the extreme (such as top versus bottom 10% or top versus bottom 20% and so on, Supplementary Fig. S5).

To compare the GI-based survival prediction with the individual gene approach, we implemented an analogous scheme for individual genes where the gene expression values were discretized into 3 expression levels (low, medium and high), and the discretized representation was used as a predictor variable in a controlled Cox regression model to obtain the significance (p-value) of each gene with respect to survival prediction. The most significant predictors (top 5%) were chosen and precisely used as described for the GIs survival prediction procedure. An analogous procedure was used to estimate the prediction accuracy based on both individual gene effects and GIs.

### Identifying gene target(s)-specific GIs

To investigate the GIs involving specific genes of interest (e.g., one or more target genes inhibited by a drug), we used a modified gene set-specific FDR approach. For a set of one or more genes X, we compare the GI significance (Cox regression p-value) of GIs involving any gene in X across the quadruples derived from step 2 (Molecular enrichment/depletion). We defined significant GI interactions as those where the GI significance is more extreme than (lower p-value, higher quantile) the 90^th^ quantile of shuffled GIs involving any member of X.

### Characterization of differential GI activation between drug-response groups

We retrieved the drug response data as explained in the first subsection of Methods; some patients have response information, and some do not. We inferred the GIs involving each drug’s known target gene based on the TCGA samples that do not have that drug’s response information to avoid circularity and filtered based on FDR restricted to the target-specific GIs (using target-specific FDR above). For each of the drug-specific GIs, we compared its activation frequency (whether the GI was functionally active or inactive) among the responders (stable disease, partial response and complete response categories) and non-responders group (clinical progressive disease categories), using one-sided Fisher’s exact test (Fisher, 1922), where the alternative hypothesis was that negative (respectively, positive) GIs are more frequently active among responders (respectively, non-responders). However, given the extremely small and imbalanced sample sizes, and the conservative nature of Fisher’s exact test (Berkson, 1978), we tested whether the overall distribution of the obtained odds-ratios are lower than those obtained using randomly shuffled drug-response labels, using one-sided Wilcoxon tests (Wilcoxon, 1945). We thus obtained a p-value for each drug-cancer type pair, segregated by GI type.

Then, for each gene pair in the inferred GIs list and, as a control, in a shuffled list of size 10x as the original GIs list size, we computed the distance between the genes in the PPI network (Schaefer et al., 2012). We then used one-sided Wilcoxon tests (Wilcoxon, 1945) to assess whether GI gene pairs are closer to each other than random expectation. Alternatively, we also compare the number of directly GI gene pairs having direct interaction using one-sided Fisher’s exact test (Fisher, 1922).

### Characterization of tissue-specific effect of cancer driver genes

A study (Rubio-Perez et al., 2015) of genes’ somatic mutation profile across cancer types has identified and characterized the tissue specificity of 459 candidate drivers. For each such candidate, we matched the driver role annotation (oncogene or tumor-suppressor) obtained from the Cosmic Census (Futreal et al., 2004) cancer genes dataset, to obtain a set of 33 tumor suppressors and 25 oncogenes matching the tissue/tumor type annotations. For each of these 58 genes, we calculated the significant GI interactions involving this gene (target-specific FDR). We further excluded genes with 5 or fewer interactions or with 300 or fewer samples where they are expressed, reducing the set of genes of interest to 20 tumor suppressors and 15 oncogenes spanning 10 cancer types (Supplementary Table S3). For a gene, a sample-specific risk score was calculated based on the functionally active GI partners of the gene (as above for the drug response analysis above), but only considering high activity bin for oncogenes and low-activity bins for tumor suppressors. For each gene, the cancer types are partitioned into affected types (cancer types affected by the driver) and the other unaffected cancer types. Using a one-sided Wilcoxon rank-sum test, we tested for higher risk score in the samples in the affected cancer type in comparison to those in unaffected types. After correcting for multiple hypotheses testing, 15 out of the 35 (~43%) driver genes were found to have significant tissue-specific GI-based risk score (FDR q-value < 0.1, Supplementary Table S3).

### Breast cancer tumor stratification

We represent a tumor sample as a vector indicating the functional activity status of each predicted GI. This provides a survival-cognizant alternative to the widely-used gene expression profile representation of a sample. We used this representation to partition the METABRIC (as well as independently for TCGA) breast cancer patients into clusters using Non-Negative Matrix Factorization (NMF using the brunet algorithm and assigning each sample to the cluster with the highest weight) (Lee and Seung, 2000; Paatero and Tapper, 1994), which has suitable statistical properties and has been shown to be effective in a variety of contexts (Lee and Seung, 1999). Since NMF requires a predetermined number of clusters, we performed the analysis for 2-15 clusters, and assessed their fitness using Dunn’s index (Dunn†, 1974), which quantifies compactness within and separation across clusters. The hazard-ratio significance values were computed for each pair of clusters, while the p-values were generated using multi-class log-rank test. For comparison purposes, to match our estimated clusters’ sizes to previously published METABRIC cluster sizes (~900 samples), we constrained the number of samples in each cluster to the 1000 samples that were found to be most highly associated with the cluster. Each cluster’s GI profiles were constructed as follows. Our clustering approach – NMF, assigned each GI to a single cluster. For each cluster, and for each of the 12 GI-types (6 bins in Fig. 1 and the two directional effects on survival), we obtain the frequency of GIs of that type, relative to all GI assigned to the clusters.

Mutation frequency analysis was performed on the TCGA clusters. We defined the gene-wise mutation frequency as the fraction of samples in the cluster in which the gene has a mutation predicted to be deleterious as explained in the next paragraph. Then, we tested whether the mutation frequency distribution of each gene differs significantly across clusters using Chi-square tests. The genes with significant Chi-square statistic (FDR q-value < 0.1) were then used to illustrate the mutational profiles of the clusters.

Each mutation’s predicted effect on the protein function was obtained from the cBioPortal repository. Out of the 196 differentially mutated genes, 138 genes had matching extended mutation information indicating their SIFT (sorts intolerant from tolerant amino acid substitutions) and PolyPhen (polymorphism phenotyping) predictions. We calculated a gene-wise fraction of mutations predicted to have a significant effect on the protein, separately for SIFT and PolyPhen.

The breast cancer subtypes were derived using the widely accepted PAM50 algorithm (Bernard et al., 2009). The METABRIC PAM50 subtypes were annotated in the original publication (Curtis et al., 2012), while the TCGA breast cancer subtypes were calculated using the original published class centroids (Bernard et al., 2009).

### Software availability

The EnGIne software is available on GitHub [https://github.com/asmagen/EncyclopediaGeneticInteractions].

The GI network is accessible online via a web portal [https://amagen.shinyapps.io/cancerapp/].

## Acknowledgement

This research is supported in part by the Intramural Research Program of the National Institutes of Health, National Cancer Institute. S.H. is funded in part by NSF award 1564785.

## References

Akaogi, K., Nakajima, Y., Ito, I., Kawasaki, S., Oie, S., Murayama, A., Kimura, K., and Yanagisawa, J. (2009). KLF4 suppresses estrogen-dependent breast cancer growth by inhibiting the transcriptional activity of ERalpha. Oncogene 28, 2894–2902.

Ashworth, A., Lord, C.J., and Reis-Filho, J.S. (2011). Genetic interactions in cancer progression and treatment. Cell 145, 30–38.

Benhamou, S., and Sarasin, A. (2002). ERCC2/XPD gene polymorphisms and cancer risk. Mutagenesis 17, 463–469.

Berkson, J. (1978). In dispraise of the exact test. Do the marginal totals of the 2X2 table contain relevant information respecting the table proportions? J. Stat. Plan. Inference 2, 27–42.

Bernard, P.S., Parker, J.S., Mullins, M., Cheung, M.C.U., Leung, S., Voduc, D., Vickery, T., Davies, S., Fauron, C., He, X., et al. (2009). Supervised risk predictor of breast cancer based on intrinsic subtypes. J. Clin. Oncol. 27, 1160–1167.

Bernard-Gallon, D., Bosviel, R., Delort, L., Fontana, L., Chamoux, A., Rabiau, N., Kwiatkowski, F., Chalabi, N., Satih, S., and Bignon, Y.-J. (2008). DNA repair gene ERCC2 polymorphisms and associations with breast and ovarian cancer risk. Mol. Cancer 7, 36.

Bommi-Reddy, A., Almeciga, I., Sawyer, J., Geisen, C., Li, W., Harlow, E., Kaelin Jr., W.G., and Grueneberg, D.A. (2008). Kinase requirements in human cells: III. Altered kinase requirements in VHL-/- cancer cells detected in a pilot synthetic lethal screen. Proc. Natl. Acad. Sci. U. S. A. 105, 16484–16489.

Brough, R., Frankum, J.R., Costa-Cabral, S., Lord, C.J., and Ashworth, A. (2011). Searching for synthetic lethality in cancer. Curr. Opin. Genet. Dev. 21, 34–41.

Chang, K., Creighton, C.J., Davis, C., Donehower, L., Drummond, J., Wheeler, D., Ally, A., Balasundaram, M., Birol, I., Butterfield, Y.S.N., et al. (2013). The Cancer Genome Atlas Pan-Cancer analysis project. Nat. Genet. 45, 1113–1120.

Chen, R., Valencia, I., Zhong, F., McColl, K.S., Roderick, H.L., Bootman, M.D., Berridge, M.J., Conway, S.J., Holmes, A.B., Mignery, G.A., et al. (2004). Bcl-2 functionally interacts with inositol 1,4,5-trisphosphate receptors to regulate calcium release from the ER in response to inositol 1,4,5-trisphosphate. J. Cell Biol. 166, 193–203.

Curtis, C., Shah, S.P., Chin, S.-F., Turashvili, G., Rueda, O.M., Dunning, M.J., Speed, D., Lynch, A.G., Samarajiwa, S., Yuan, Y., et al. (2012). The genomic and transcriptomic architecture of 2,000 breast tumours reveals novel subgroups. Nature 486, 346–352.

Dunn†, J.C. (1974). Well-Separated Clusters and Optimal Fuzzy Partitions. J. Cybern. 4, 95–104.

Eisenhauer, E.A., Therasse, P., Bogaerts, J., Schwartz, L.H., Sargent, D., Ford, R., Dancey, J., Arbuck, S., Gwyther, S., Mooney, M., et al. (2009). New response evaluation criteria in solid tumours: Revised RECIST guideline (version 1.1). Eur. J. Cancer 45, 228–247.

Fisher, R.A. (1922). On the Interpretation of χ 2 from Contingency Tables, and the Calculation of P. J. R. Stat. Soc. 85, 87.

Fong, C.Y., Gilan, O., Lam, E.Y.N., Rubin, A.F., Ftouni, S., Tyler, D., Stanley, K., Sinha, D., Yeh, P., Morison, J., et al. (2015). BET inhibitor resistance emerges from leukaemia stem cells. Nature 525, 538–+.

Futreal, P.A., Coin, L., Marshall, M., Down, T., Hubbard, T., Wooster, R., Rahman, N., and Stratton, M.R. (2004). A census of human cancer genes. Nat. Rev. Cancer 4, 177–183.

Gonzalez-Perez, A., Perez-Llamas, C., Deu-Pons, J., Tamborero, D., Schroeder, M.P., Jene-Sanz, A., Santos, A., and Lopez-Bigas, N. (2013). IntOGen-mutations identifies cancer drivers across tumor types. Nat. Methods 10, 1081–1082.

Harrell, F.E., Lee, K.L., and Mark, D.B. (2005). Prognostic/Clinical Prediction Models: Multivariable Prognostic Models: Issues in Developing Models, Evaluating Assumptions and Adequacy, and Measuring and Reducing Errors. In Tutorials in Biostatistics, Statistical Methods in Clinical Studies, pp. 223–249.

Harrington, D.P., and Fleming, T.R. (1982). A class of rank test procedures for censored survival data. Biometrika 69, 553–566.

Hartwell, L.H., Szankasi, P., Roberts, C.J., Murray, A.W., and Friend, S.H. (1997). Integrating genetic approaches into the discovery of anticancer drugs. Science (80-. ). 278, 1064–1068.

Jerby, L., Waldman, Y., Weinstock, A., Geiger, T., and Ruppin, E. Genome-wide detection of synthetically-lethal genes uncovers a novel repository of selective cancer targets. 1–9.

Jerby-Arnon, L., Pfetzer, N., Waldman, Y.Y., McGarry, L., James, D., Shanks, E., Seashore-Ludlow, B., Weinstock, A., Geiger, T., Clemons, P.A., et al. (2014). Predicting cancer-specific vulnerability via data-driven detection of synthetic lethality. Cell 158, 1199–1209.

Kaelin, W.G. (2005). The concept of synthetic lethality in the context of anticancer therapy. Nat. Rev. Cancer 5, 689–698.

Kawai, T., Forrester, S.J., O’Brien, S., Baggett, A., Rizzo, V., and Eguchi, S. (2017). AT1 receptor signaling pathways in the cardiovascular system. Pharmacol. Res. 125, 4–13.

Kelley, R., and Ideker, T. (2005). Systematic interpretation of genetic interactions using protein networks. Nat. Biotechnol. 23, 561–566.

Kroll, E.S., Hyland, K.M., Hieter, P., and Li, J.J. (1996). Establishing genetic interactions by a synthetic dosage lethality phenotype. Genetics 143, 95–102.

Lambert, M., Jambon, S., Depauw, S., and David-Cordonnier, M.H. (2018). Targeting transcription factors for cancer treatment. Molecules 23.

Law, V., Knox, C., Djoumbou, Y., Jewison, T., Guo, A.C., Liu, Y., MacIejewski, A., Arndt, D., Wilson, M., Neveu, V., et al. (2014). DrugBank 4.0: Shedding new light on drug metabolism. Nucleic Acids Res. 42.

Lee, D.D., and Seung, H.S. (1999). Learning the parts of objects by non-negative matrix factorization. Nature 401, 788–791.

Lee, D.D., and Seung, H.S. (2000). Algorithms for Non-negative Matrix Factorization. In NIPS, pp. 556–562.

Lee, J.S., Das, A., Jerby-Arnon, L., Arafeh, R., Auslander, N., Davidson, M., McGarry, L., James, D., Amzallag, A., Park, S.G., et al. (2018). Harnessing synthetic lethality to predict the response to cancer treatment. Nat. Commun. 9.

Li, H., Fang, Y., Niu, C., Cao, H., Mi, T., Zhu, H., Yuan, J., and Zhu, J. (2018). Inhibition of cIAP1 as a strategy for targeting c-MYC–driven oncogenic activity. Proc. Natl. Acad. Sci. 115, E9317–E9324.

Lord, C.J., McDonald, S., Swift, S., Turner, N.C., and Ashworth, A. (2010). A high-throughput RNA interference screen for DNA repair determinants of PARP inhibitor sensitivity. DNA Repair (Amst). 7, 2010–2019.

Lu, X., Kensche, P.R., Huynen, M. a, and Notebaart, R. a (2013). Genome evolution predicts genetic interactions in protein complexes and reveals cancer drug targets. Nat. Commun. 4, 2124.

Luo, J., Solimini, N.L., and Elledge, S.J. (2009a). Principles of Cancer Therapy: Oncogene and Non-oncogene Addiction. Cell 136, 823–837.

Luo, J., Emanuele, M.J., Li, D., Creighton, C.J., Schlabach, M.R., Westbrook, T.F., Wong, K.K., and Elledge, S.J. (2009b). A Genome-wide RNAi Screen Identifies Multiple Synthetic Lethal Interactions with the Ras Oncogene. Cell 137, 835–848.

Martin, S.A., McCarthy, A., Barber, L.J., Burgess, D.J., Parry, S., Lord, C.J., and Ashworth, A. (2009). Methotrexate induces oxidative DNA damage and is selectively lethal to tumour cells with defects in the DNA mismatch repair gene MSH2. EMBO Mol. Med. 1, 323–337.

Matsushita, K., Shimada, H., Ueda, Y., Inoue, M., Hasegawa, M., Tomonaga, T., Matsubara, H., and Nomura, F. (2014). Non-transmissible Sendai virus vector encoding c-myc suppressor FBP-interacting repressor for cancer therapy. World J. Gastroenterol. 20, 4316–4328.

McLornan, D.P., List, A., and Mufti, G.J. (2014). Applying Synthetic Lethality for the Selective Targeting of Cancer. N. Engl. J. Med. 371, 1725–1735.

Megchelenbrink, W., Katzir, R., Lu, X., Ruppin, E., and Notebaart, R. a (2015). Synthetic dosage lethality in the human metabolic network is highly predictive of tumor growth and cancer patient survival. Proc. Natl. Acad. Sci. U. S. A. 112, 12217–12222.

Miyamoto, D.T., Zheng, Y., Wittner, B.S., Lee, R.J., Zhu, H., Broderick, K.T., Desai, R., Fox, D.B., Brannigan, B.W., Trautwein, J., et al. (2015). RNA-Seq of single prostate CTCs implicates noncanonical Wnt signaling in antiandrogen resistance. Science 349, 1351–1356.

Oakes, S.A., Scorrano, L., Opferman, J.T., Bassik, M.C., Nishino, M., Pozzan, T., and Korsmeyer, S.J. (2005). Proapoptotic BAX and BAK regulate the type 1 inositol trisphosphate receptor and calcium leak from the endoplasmic reticulum. Proc. Natl. Acad. Sci. 102, 105–110.

Paatero, P., and Tapper, U. (1994). Positive Matrix Factorization - A Nonnegative Factor Model With Optimal Utilization of Error-Estimates of Data Values. Environmetrics 5, 111–126.

Rahmutulla, B., Matsushita, K., Satoh, M., Seimiya, M., Tsuchida, S., Kubo, S., Shimada, H., Ohtsuka, M., Miyazaki, M., and Nomura, F. (2013). Alternative splicing of FBP-interacting repressor coordinates c-Myc, P27Kip1/cyclinE and Ku86/XRCC5 expression as a molecular sensor for bleomycin-induced DNA damage pathway. Oncotarget 5, 2404–2417.

Rathert, P., Roth, M., Neumann, T., Muerdter, F., Roe, J.-S.J.-S., Muhar, M., Deswal, S., Cerny-Reiterer, S., Peter, B., Jude, J., et al. (2015). Transcriptional plasticity promotes primary and acquired resistance to BET inhibition. Nature 525, 543–547.

Rong, Y.-P., Bultynck, G., Aromolaran, A.S., Zhong, F., Parys, J.B., De Smedt, H., Mignery, G.A., Roderick, H.L., Bootman, M.D., and Distelhorst, C.W. (2009). The BH4 domain of Bcl-2 inhibits ER calcium release and apoptosis by binding the regulatory and coupling domain of the IP3 receptor. Proc. Natl. Acad. Sci. 106, 14397–14402.

Rubio-Perez, C., Tamborero, D., Schroeder, M.P., Antolín, A.A., Deu-Pons, J., Perez-Llamas, C., Mestres, J., Gonzalez-Perez, A., and Lopez-Bigas, N. (2015). In Silico Prescription of Anticancer Drugs to Cohorts of 28 Tumor Types Reveals Targeting Opportunities. Cancer Cell 27, 382–396.

Ruvolo, P.P., Deng, X., and May, W.S. (2001). Phosphorylation of Bcl2 and regulation of apoptosis. Leuk. Off. J. Leuk. Soc. Am. Leuk. Res. Fund, U.K 15, 515–522.

Sajesh, B. V., Guppy, B.J., and McManus, K.J. (2013). Synthetic genetic targeting of genome instability in cancer. Cancers (Basel). 5, 739–761.

Schaefer, M.H., Fontaine, J.F., Vinayagam, A., Porras, P., Wanker, E.E., and Andrade-Navarro, M.A. (2012). Hippie: Integrating protein interaction networks with experiment based quality scores. PLoS One 7.

Steckel, M., Molina-Arcas, M., Weigelt, B., Marani, M., Warne, P.H., Kuznetsov, H., Kelly, G., Saunders, B., Howell, M., Downward, J., et al. (2012). Determination of synthetic lethal interactions in KRAS oncogene-dependent cancer cells reveals novel therapeutic targeting strategies. Cell Res. 22, 1227–1245.

Stuhlmiller, T.J., Miller, S.M., Zawistowski, J.S., Nakamura, K., Beltran, A.S., Duncan, J.S., Angus, S.P., Collins, K.A.L., Granger, D.A., Reuther, R.A., et al. (2015). Inhibition of lapatinib-induced kinome reprogramming in ERBB2-positive breast cancer by targeting BET family bromodomains. Cell Rep. 11, 390–404.

Szappanos, B., Kovács, K., Szamecz, B., Honti, F., Costanzo, M., Baryshnikova, A., Gelius-Dietrich, G., Lercher, M.J., Jelasity, M., Myers, C.L., et al. (2011). An integrated approach to characterize genetic interaction networks in yeast metabolism. Nat. Genet. 43, 656–662.

Szczurek, E., Misra, N., and Vingron, M. (2013). Synthetic sickness or lethality points at candidate combination therapy targets in glioblastoma. Int. J. Cancer 133, 2123–2132.

Szklarczyk, D., Franceschini, A., Wyder, S., Forslund, K., Heller, D., Huerta-Cepas, J., Simonovic, M., Roth, A., Santos, A., Tsafou, K.P., et al. (2015). STRING v10: Protein-protein interaction networks, integrated over the tree of life. Nucleic Acids Res. 43, D447–D452.

Turner, N.C., Lord, C.J., Iorns, E., Brough, R., Swift, S., Elliott, R., Rayter, S., Tutt, A.N., and Ashworth, A. (2008). A synthetic lethal siRNA screen identifying genes mediating sensitivity to a PARP inhibitor. EMBO J. 27, 1368–1377.

Wilcoxon, F. (1945). Individual Comparisons by Ranking Methods. Biometrics Bull. 1, 80.

Williams, C.S., Watson, A.J.M., Sheng, H., Helou, R., Shao, J., and DuBois, R.N. (2000). Celecoxib prevents tumor growth in vivo without toxicity to normal gut: Lack of correlation between in vitro and in vivo models. Cancer Res. 60, 6045–6051.

Wong, S.L., Zhang, L. V, Tong, A.H.Y., Li, Z., Goldberg, D.S., King, O.D., Lesage, G., Vidal, M., Andrews, B., Bussey, H., et al. (2004). Combining biological networks to predict genetic interactions. Proc. Natl. Acad. Sci. U. S. A. 101, 15682–15687.

Yoo, A.B., Jette, M.A., and Grondona, M. (2003). SLURM: Simple Linux Utility for Resource Management. In Job Scheduling Strategies for Parallel Processing, D. Feitelson, L. Rudolph, and U. Schwiegelshohn, eds. (Berlin, Heidelberg: Springer Berlin Heidelberg), pp. 44–60.

Yuan, Y., Allen, E.M. Van, Omberg, L., Wagle, N., Amin-Mansour, A., Sokolov, A., Byers, L. a, Xu, Y., Hess, K.R., Diao, L., et al. (2014). Assessing the clinical utility of cancer genomic and proteomic data across tumor types. Nat. Biotechnol. 32, 644–652.

Zhong, W., and Sternberg, P.W. (2006). Genome-wide prediction of C. elegans genetic interactions. Science 311, 1481–1484.

