## Supplementary Text and Figures for "Beyond synthetic lethality: charting the landscape of clinically relevant genetic interactions in cancer"

**Supplementary Results**

**GI based prediction of patient survival**

Recall that a GI is represented by a quadruple comprising of a gene pair (x,y), a symmetric bin b (1, 2, 3, 5, 6, 9; Fig. 1), and its effect on tumor fitness (positive or negative), and a GI is deemed *functionally active* in a specific tumor sample if the joint expression levels of the genes x, y, in the tumor fall in bin b. We turned to assess the extent to which *the aggregate effect* of functionally active GIs in a tumor predicts patient survival (the individual GIs are indeed inferred while considering survival but here, we are interested in the predictive power of their combined effects). The naïve tumor-specific GI survival score is computed as the difference between the number of functionally active GIs in a tumor with positive effect on survival and the number of those with negative effect on survival. We compared the GI-based survival prediction accuracies with a comparable gene-level approach (which is based on assessing the expression levels of the genes that are significantly associated with patient survival) as well as a combined gene and GI-based approach (see Methods). Fig. S4A shows the survival prediction accuracy using the widely used C-index (CI) metric, both via cross-validation within TCGA (Chang et al., 2013) and on independent METABRIC breast cancer dataset (Curtis et al., 2012) (see Methods). As shown, pan-cancer risk prediction based on the predicted GIs compares favorably with the comparable gene-level approach both in cross-validation, and more so on the METABRIC independent dataset － suggesting good generalizability. Applying the fully supervised individual gene-level approach (Huang et al., 2016) (individual gene feature selection based on association with survival, further dimensionality reduction with LASSO Cox and a final Cox model to obtain genes’ coefficients for survival risk prediction) yields a comparable accuracy in cross validation (CI ≈ 0.63) but a lower accuracy over the independent dataset (CI ≈ 0.55). Finally, a previous study has reported a CI of 0.71 using a supervised approach specifically on Kidney Renal Clear Cell Carcinoma (KIRC). We ensured that a lower gene-wise accuracy that we observe is not simply due to our filtering and implementation (methods); applying the fully supervised individual gene-level approach on KIRC subset of the filtered dataset yields CI ≈ 0.71, recapitulating the accuracy reported in the original publication (Huang et al., 2016).

In Fig. S4B, we present the performance of the GI-based approach using each of the 6 GI types. As evident, interactions involving both genes in their wild type mid-activity levels have negligible predictive power on survival, testifying that more extreme levels of gene expression tend to be involved in functional GIs affecting survival. Supplementary Fig. S5 shows the performance based on an alternative metric where we dichotomized the extreme (at varying thresholds from 10% to 50%) predicted low- and high-risk groups and quantified the difference in their area under their KM survival curves (see Methods). The survival prediction performance of the full compendium of 12 (positive and negative) GI types is shown in Supplementary Fig. S6.

**Potential biological significance of mutations across BRCA clusters**

Cyclin dependent kinase *CDK12*, involved in DNA damage response, is known to affect *BRCA1* transcription (Blazek et al., 2011). Likewise, cyclin *CDKN1B* is known to be mutated in luminal breast cancer (Stephens et al., 2012). Our analysis suggests a more specific enrichment of this mutation in subset of Luminal-A tumors (cluster 9 in Fig. 4B). The Her-2 breast cancer subtype is conventionally characterized by overexpression of HER2, encoded by the gene *ERBB2*, but rare activating mutations of this gene have also been implicated in the disease, independent of overexpression (Sun et al., 2015). Our analysis reveals an enrichment of *ERBB2* mutations in cluster 1, which includes only a small subset of annotated Her-2 subtype tumors. Most of the *ERBB2* missense mutations (11 out of 13) are predicted to have a significant impact on the protein function (SIFT and PolyPhen measures, Methods). This may suggest a distinct oncogenic mechanism underlying this subclass of tumors, which may express a Her-2-like subtype including a few samples where HER2 is over-expressed and many others where it is not over-expressed, but mutated. As another example, the *FCGR2B* mutation modulates Her-2 breast tumor’s response to Trastuzumab (Norton et al., 2014). Our analysis reveals a specific enrichment of *FCGR2B* mutations in cluster 6, which includes several other breast cancer subtypes besides Her-2, pointing to the potential relevance of these mutations in tumors belonging to other cancer subtypes as well (we emphasize that no mutation data was used at all in the GIs inference via EnGIne).

**Supplementary Notes**

1. We assessed the robustness of threshold selection throughout the EnGIne pipeline, including:
   a. Modifications of bin quantile boundaries: we kept the 3×3 structure and either increased or decreased the corner bins by 10%, corresponding to ~170 added or removed samples per bin
   b. Log-rank p-value based most significant quantile: scanning between top 5% – 30%
   c. Cox regression FDR threshold: scanning between 0.01 – 0.1

Notably, the relatively small changes in binning thresholds described in (a) result in moving hundreds of samples in or out of the bins (170 ~ 340 samples), thus it is a significant change of the bin size and composition. For computational tractability of exploring a large parameter space, we limited the set of genes to 557 Cosmic Cancer Census genes (Tier 1 and 2) corresponding to 154,846 possible gene pair combinations. As shown in Table S7, for the alternate binning, across all log-rank thresholds, 57%-96% of the gene pairs are recapitulated. Likewise, setting the log-rank quantile threshold to 0.8 (top 20% retained), across various Cox FDR thresholds, 45%-56% of the GI detected using the default setting are captured. For instance, at log-rank threshold = 0.8 and Cox regression FDR = 0.1 yields ~33,000 GIs (regardless of the binning threshold), and these include 53%~55% of the GIs defined using the original binning method. While the EnGIne pipeline offers users to set the above parameters, to demonstrate the utility of EnGIne in the main results we have used a more stringent Log-rank p-value based quantile and Cox FDR in the genome wide analysis to make the execution time of the pipeline tractable.

2. To assess whether the full GIs network is dominated by correlated gene expression patterns, we compared between correlations found between GI pairs to shuffled GIs. For a pair of GIs A = (x_1_,y_1_) and B = (x_2_,y_2_), we defined the Pearson correlation statistic (PCS) between A and B as $Min\left( \rho\left( x_{1},x_{2} \right),\rho(y_{1},y_{2}) \right)$, quantifying whether the two GIs are independently inferred. Then, we calculated the PCS for GI pairs within each GI type and compared them to PCS calculated for the shuffled GI pairs from the same GI type. Given the high number of GIs and the infeasible number of pairs to test, we selected up to 1000 GIs randomly from each GI type and calculated the corresponding shuffled GI list of the same length. The PCS comparisons indicated similar distributions of the actual and shuffled GIs across all GI types (Supplementary Figure S2B, Table S8). Although GI type +9 exhibited higher PCS relative to shuffled, the effect size was marginal (Table S8). Thus, we conclude that the full GI network is not dominated by correlated gene expression patterns.
3. The TCGA breast cancer cohort is composed of several subsets of independent studies, with potential batch effects and other confounders. To assess the quality of the TCGA breast cancer molecular and clinical data, we analyzed the association with survival of 79 known breast cancer genes (Intogen dataset (Gonzalez-Perez et al., 2013)). We find 31 (~40%) of the genes to be significantly associated with survival (P < 0.05). Finally, although the TCGA cohort may be noisy due to the data collection procedures, we find the survival prediction accuracy of the discovered GIs to generalize to the METABRIC dataset, supporting a reasonable quality of the data.
4. We assessed genomic-instability and tumor purity as potential confounding factors and found that the vast majority of the discovered GIs remain highly significant. The genomic instability index measures the relative amplification or deletion of genes in a tumor based on the SCNA (Bilal et al., 2013) and the tumor purity is an estimate of the proportion of cancer cells in the sample (Aran et al., 2015). We recomputed the controlled Cox survival step (Fig. 1E) for all the candidates and shuffled candidates obtained from the previous step of the pipeline, with the addition of the two covariates － genomic instability and tumor purity, to calculate the empirical FDR threshold (0.01 quantile). We obtain 77630 confounder-corrected GIs where (1) 60302 of the 71946 original GIs and (2) 1405 of the 1704 original PPI GIs were detected. Modifying the FDR threshold to 0.1 yields a list of 154450 confounder-corrected GIs which contain almost all of the 72k original GIs with the exception of 48 GIs that are not included (these 48 GIs remain the exception when increasing the FDR quantile threshold to 0.2, reflecting the set of highly confounded GIs, Supplementary Table S6). 45 of the confounded GIs involve DEFB21, a member of the beta subfamily of defensins (antimicrobial peptides), suggesting that the majority of the confounded effects are limited to one gene as oppose to a wide-spread phenomenon. None of the GIs is discussed in the main text are affected by this additional confounding control.
5. We sought to validate the discovered GIs in additional datasets but to the best of our knowledge, the TCGA and METABRIC are the only human datasets with long-term survival follow-up, clinical-demographic group annotations and sufficient (thousands) of samples for robust division into bins. Nonetheless, we validated the final list of 1704 GIs on the NKI dataset (van de Vijver et al., 2002) and found an enriched proportion of significance GIs only in bin 1 (0.125, 5 out of 40, 2.5 fold for P < 0.05). This potentially reflects the error in binning over small datasets with no complete clinical-demographic data.

**Supplementary Figures**

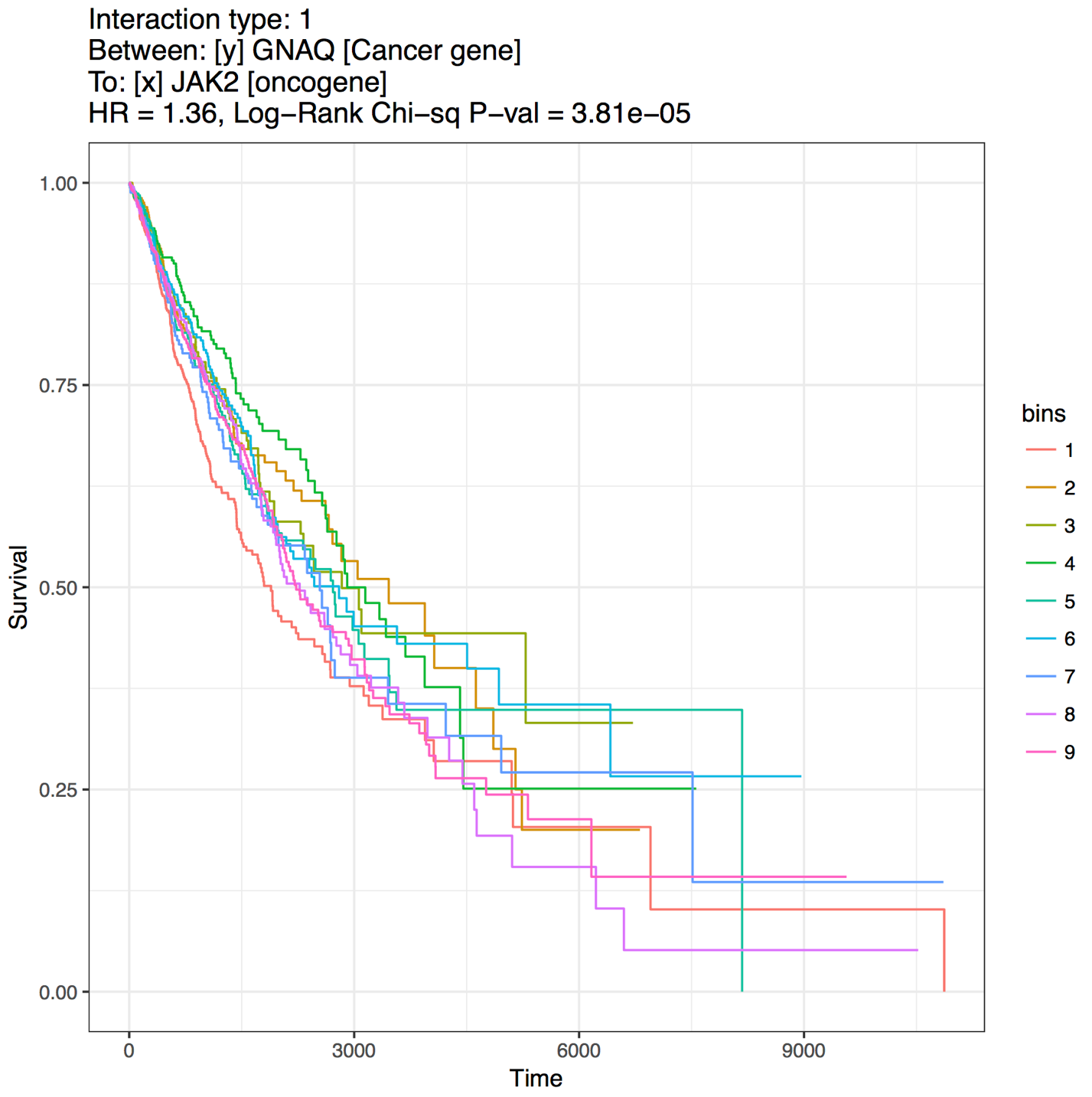

**Figure S1** - The Interaction between the cancer genes GNAQ  and JAK2  as an example of a positive interaction in bin-1.

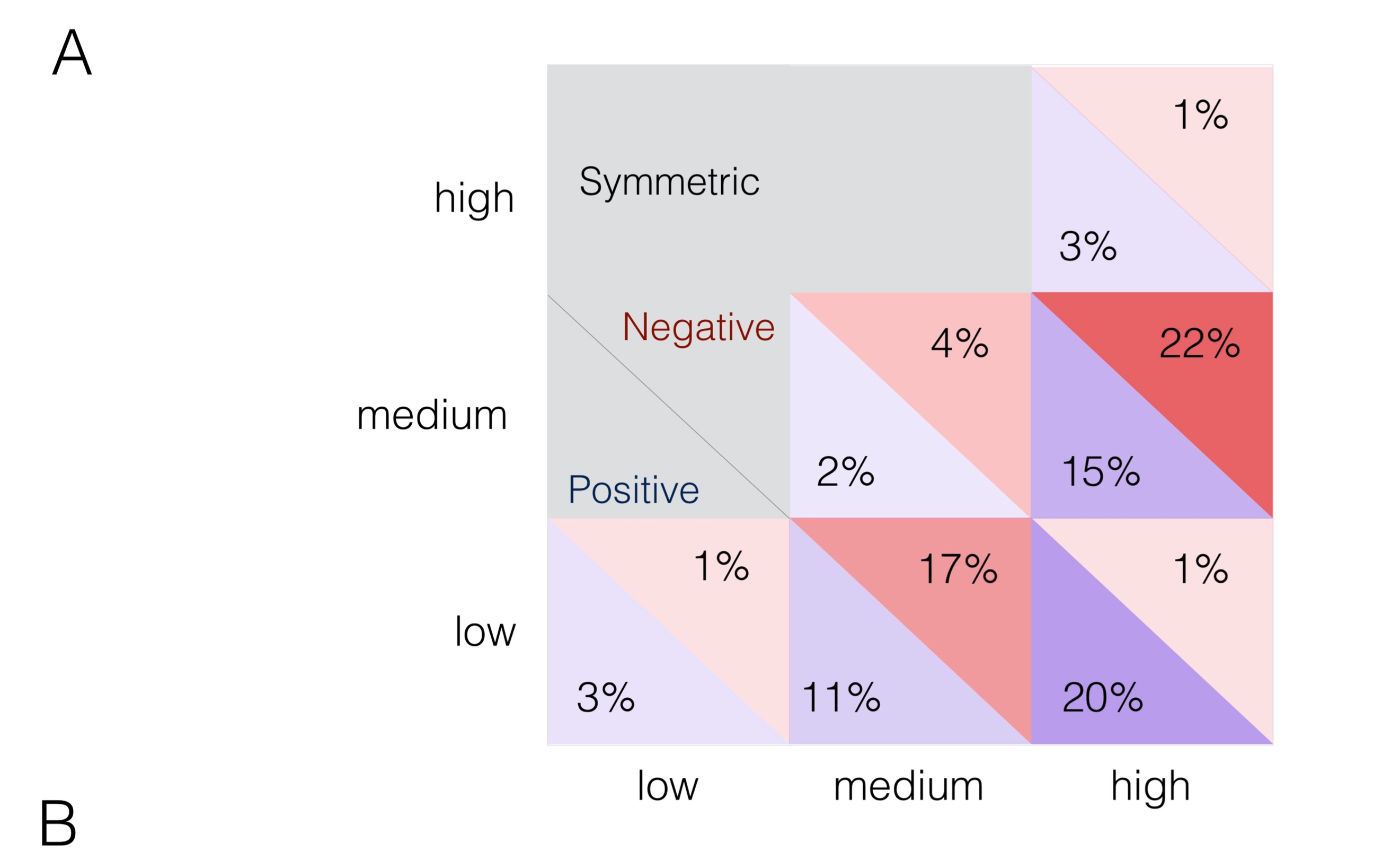

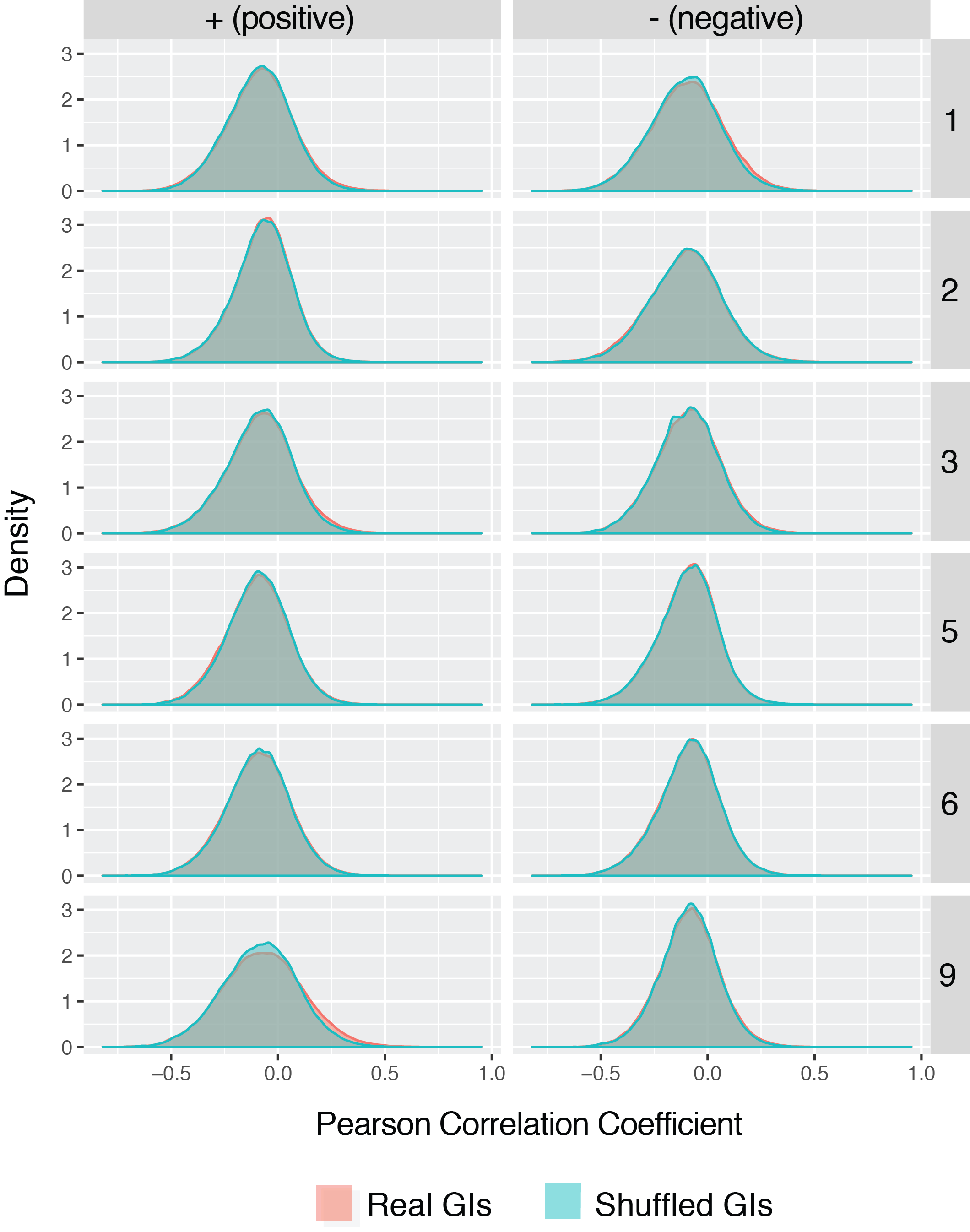

**Figure S2** – **(A) Distribution of the 71,946 significant GIs across 12 joint activity bins.** The fractions of GIs in each bin are shown for GIs with positive (blue) and negative (red) effect on tumor fitness. Only the data in the lower triangle of the matrix are shown as the GIs are symmetric relative to the genes in a pair. **(B) Correlation between GI pairs and shuffled GI pairs across the 12 activity bins.** Minimum statistic distribution of Pearson correlation between genes defining the Real GIs (non-shuffled, red) and Shuffled GIs (blue).

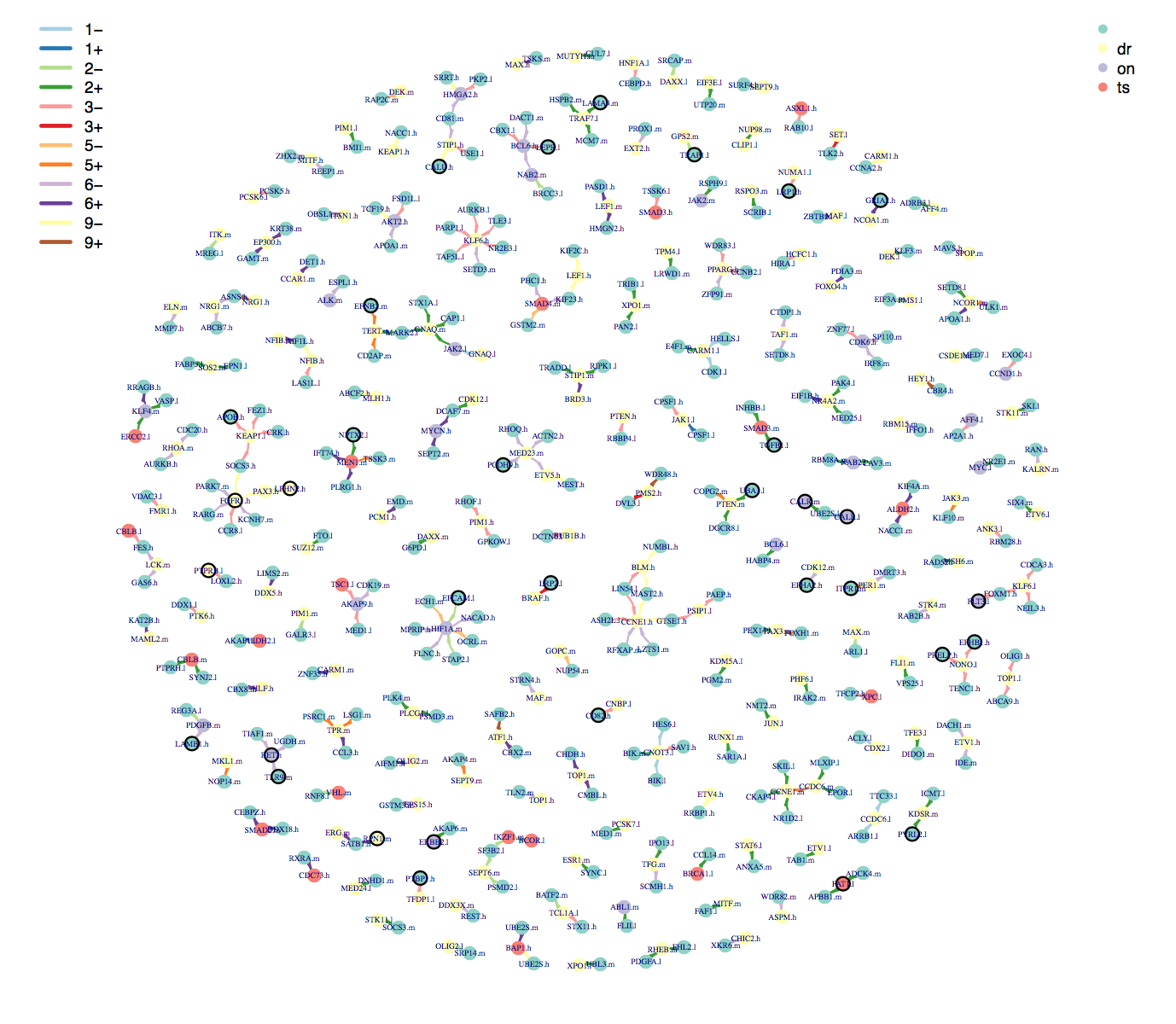
**Figure S3** - The GI-network involving the known driver genes.

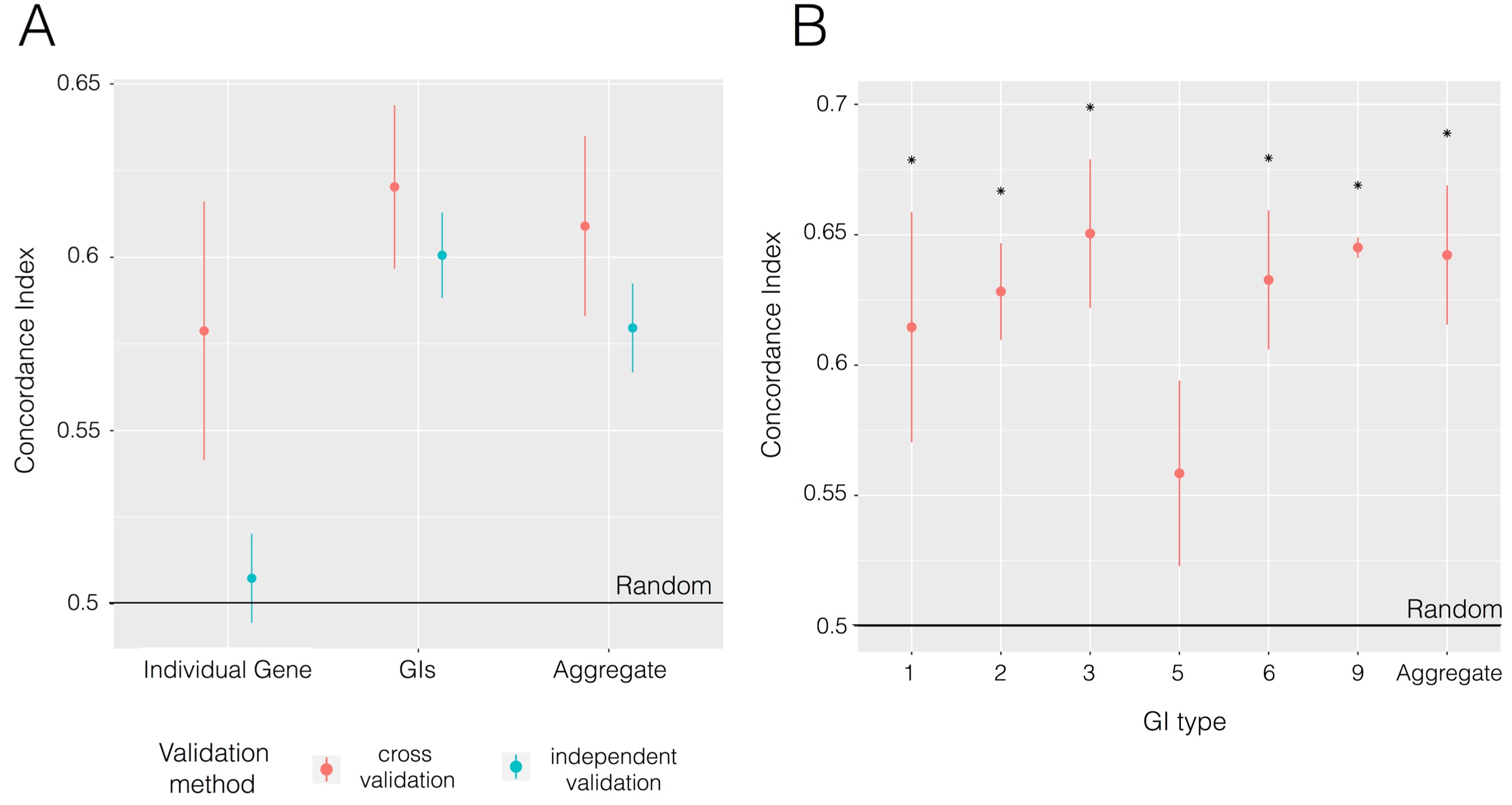

**Figure S4 – (A-B) Survival prediction accuracy.** The prediction accuracy (y-axes) is measured by the concordance of predicted and observed survival (C-index), where 0.5 represents the null expectation. (A) C-index comparison between the individual gene-wise approach, the GI-based approach and the approach that aggregates both individual gene signals and the GIs. The accuracy is estimated based on cross-validation within TCGA (red) as well as in an independent METABRIC breast cancer dataset (blue). The individual gene approach performs poorly in cross validation and fails in the independent validation while the GIs achieve superior accuracy in both validation approaches. (B) Comparison of cross-validation prediction accuracies when the 6 different GIs types are used in isolation. ‘All’ on x-axis refers to overall GI-based prediction accuracy using all six types (Significant predictions relative to null expectation (P-value < 0.01) are marked by an asterisk).

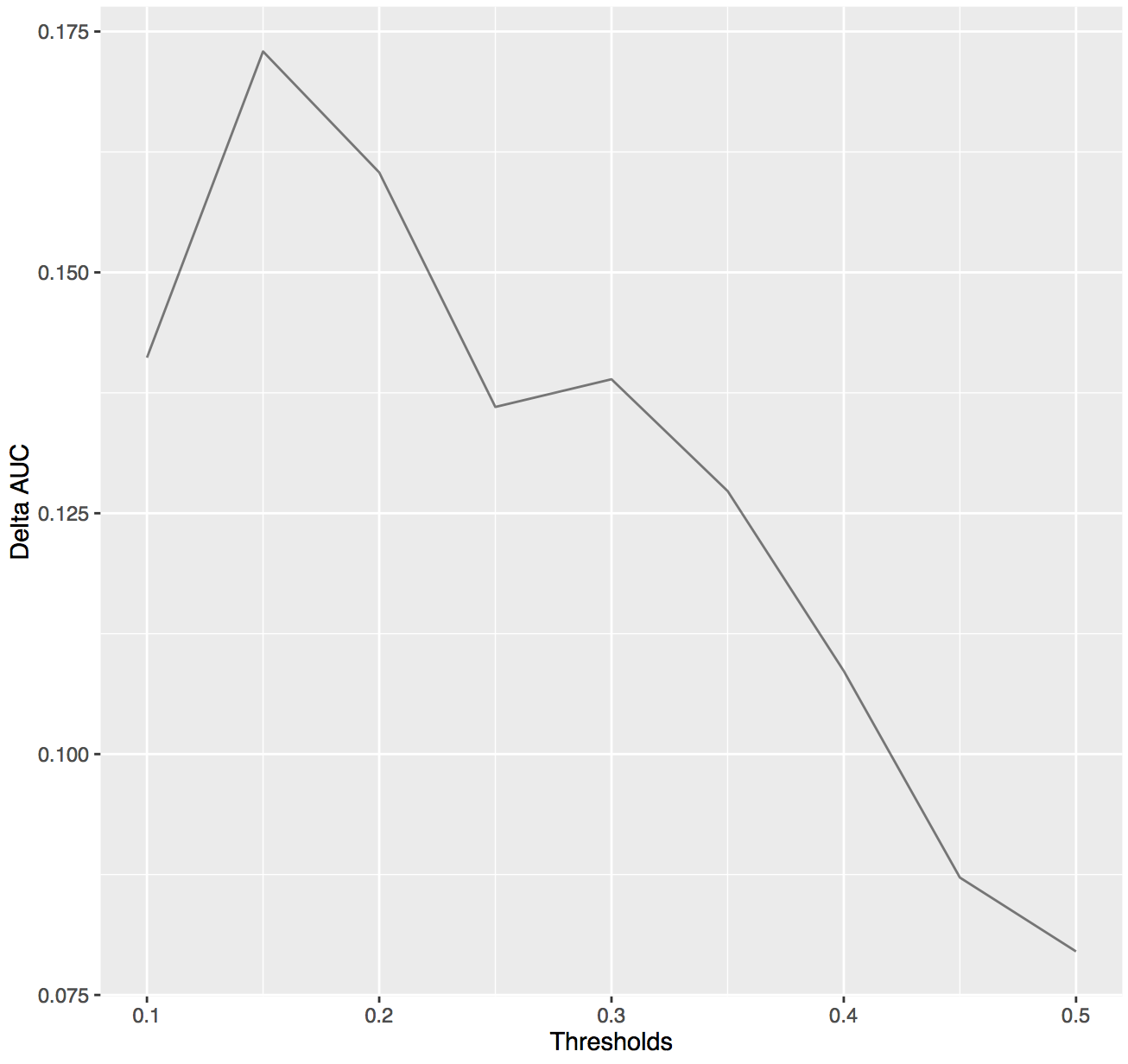

**Figure S5** – Survival analysis performance scores based on an alternative metric where we dichotomized the extreme (at varying thresholds from 10% to 50%) predicted low- and high-risk groups and quantified the difference in their area under their Kaplan Meyer (KM) survival curves.

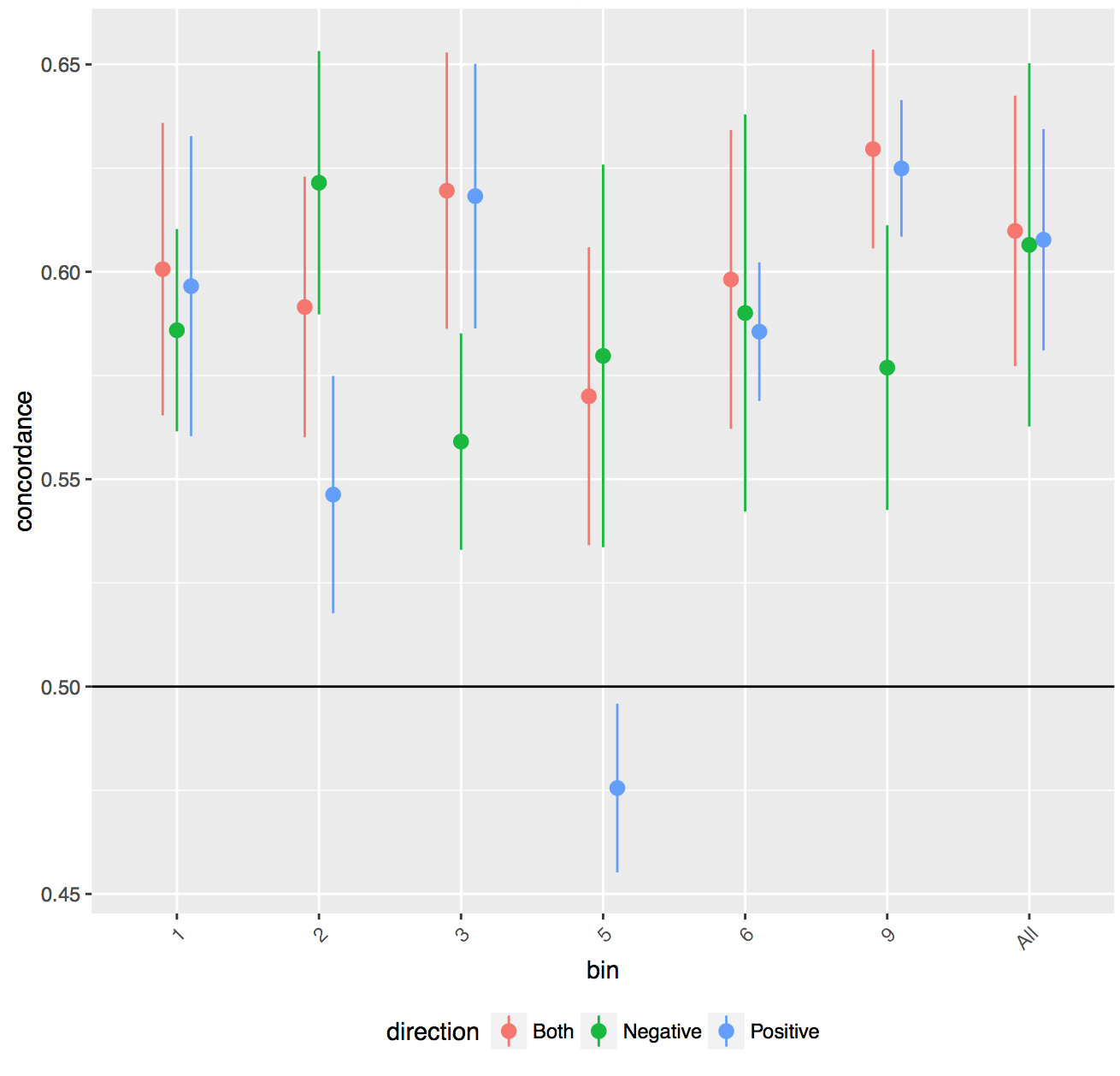
**Figure S6** – GI-based approach survival risk prediction performance restricting to GIs in each of the 6 bins and further segregated them into those with positive and negative effects on survival.

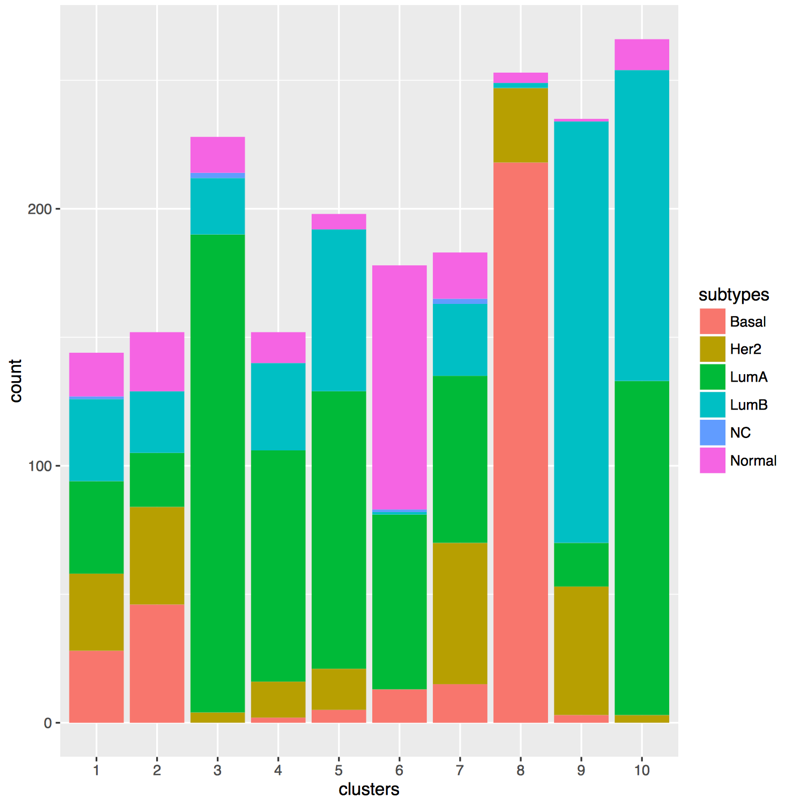

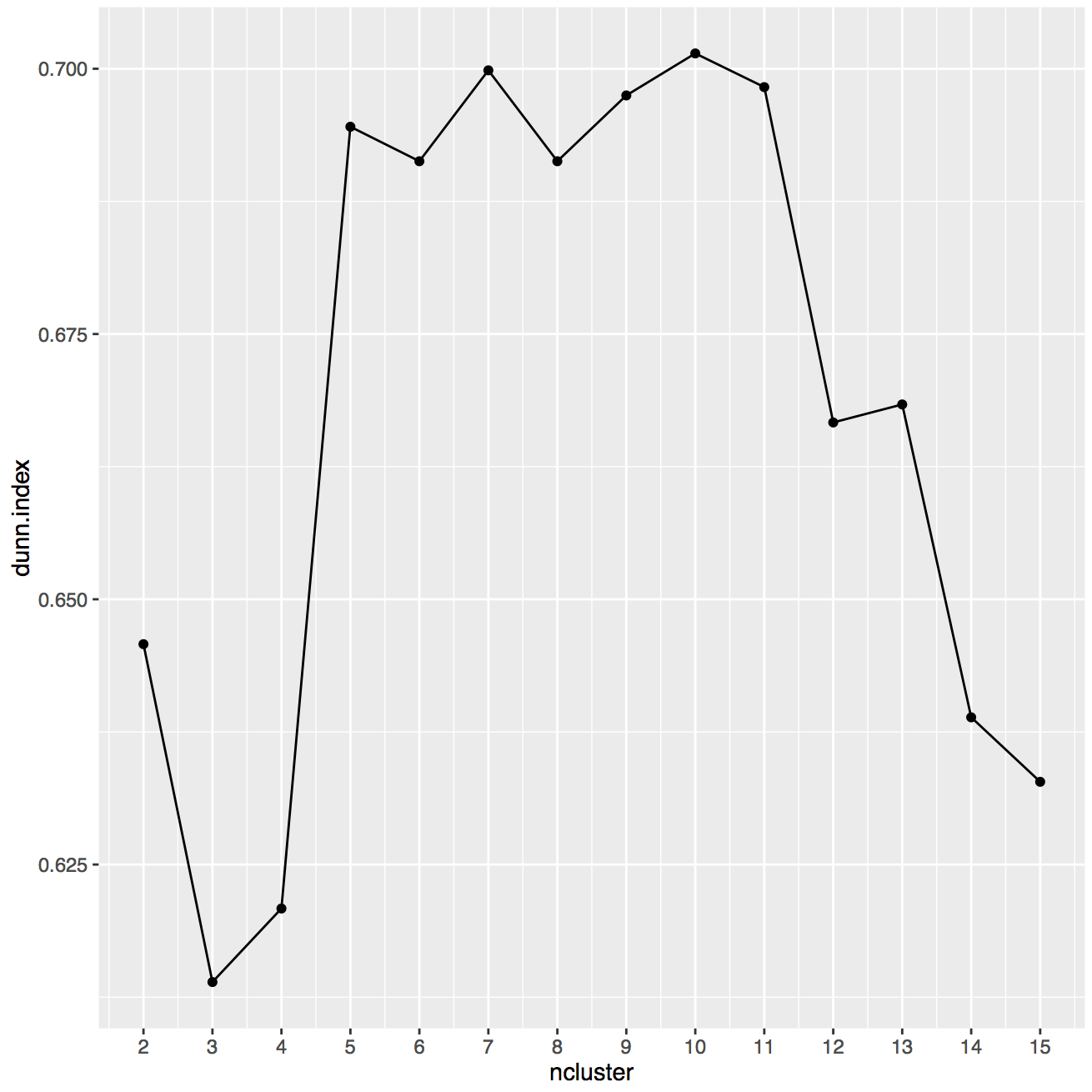

B

AC

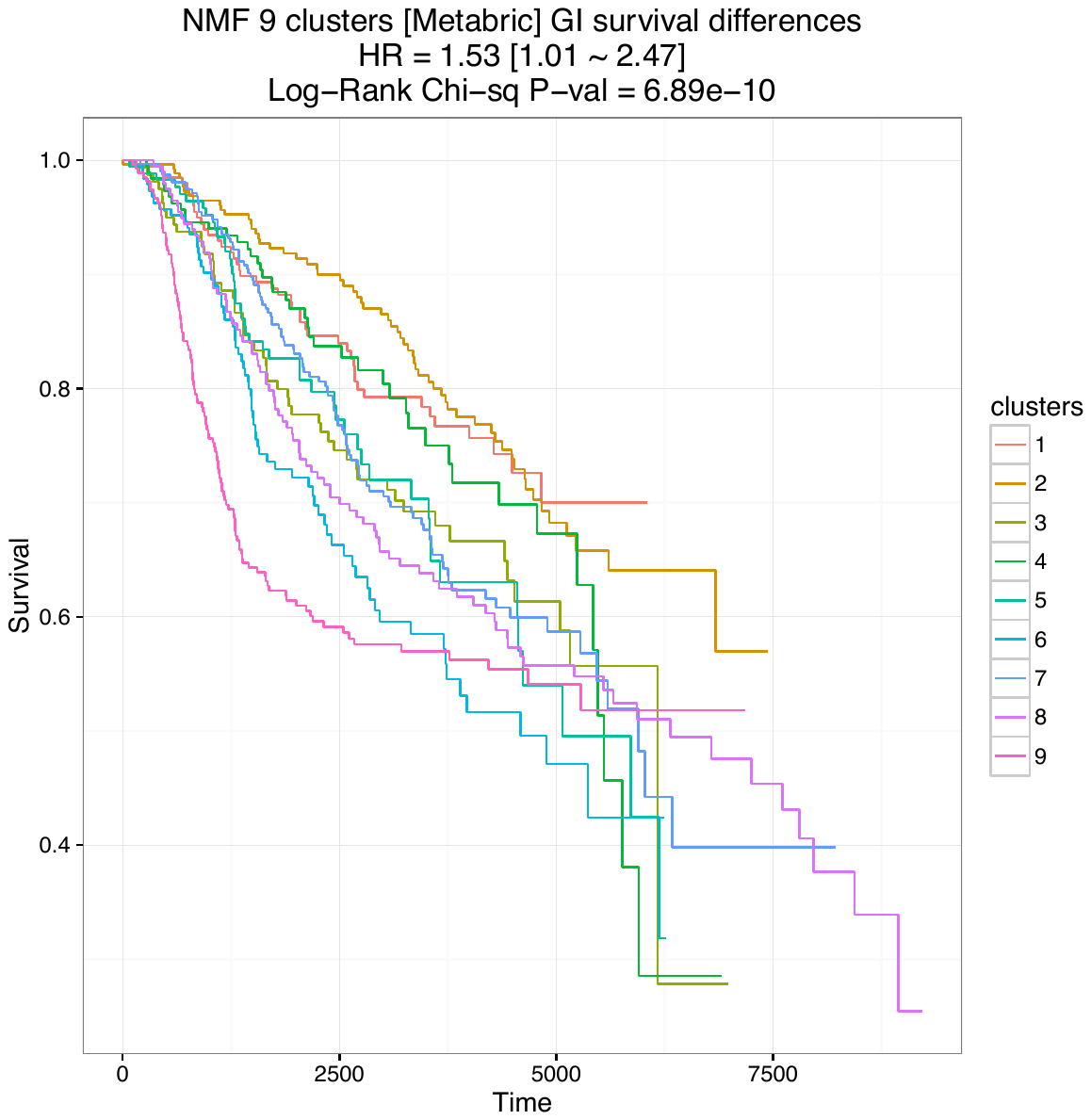

CCt

**Figure S7** – (A) Dunn’s index clustering quality score for varying number of clusters. (B) Breast cancer subtype distribution in NMF clustering using 10 clusters. (C) Survival characteristics of clustering analysis performed using the full GI network.

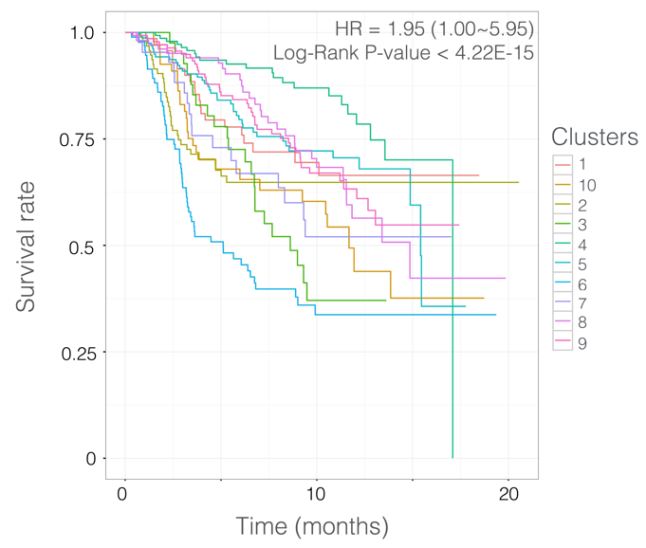

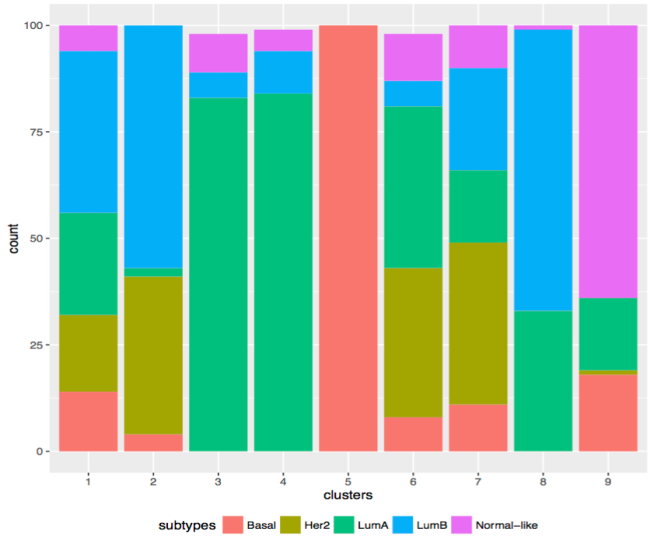

C

B

AC

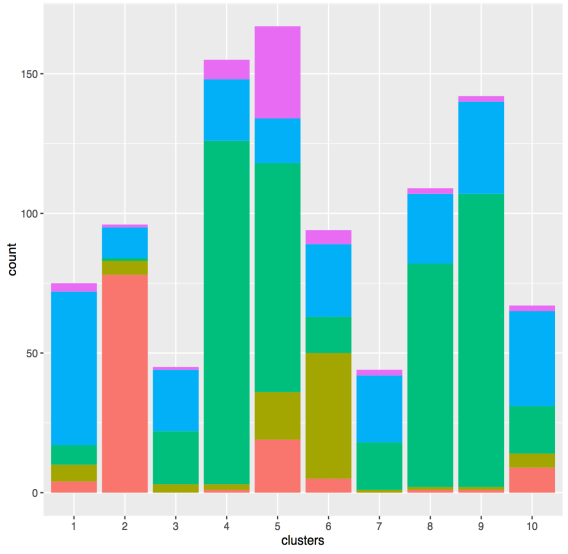

**Figure S8** –**Cluster clinical subtype composition based on PAM50 breast cancer sub-typing** (Bernard et al., 2009). (A) METABRIC original publication clustering survival trends. (B) Cluster composition based on refined GI clusters, (C) Cluster composition provided in the original METABRIC publication.

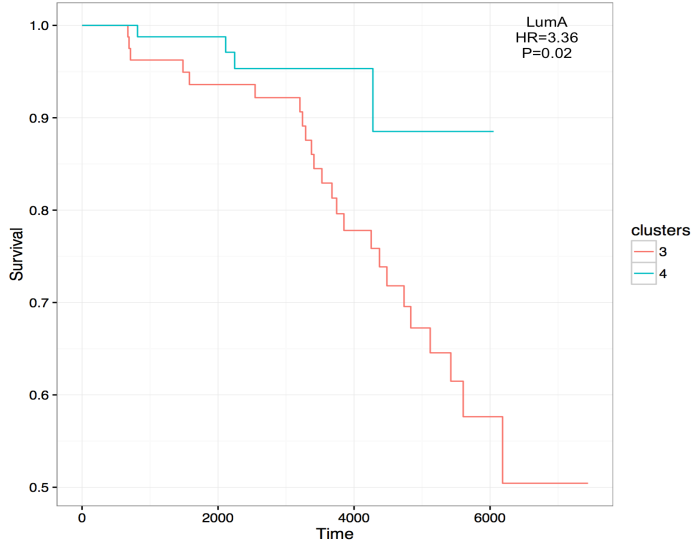

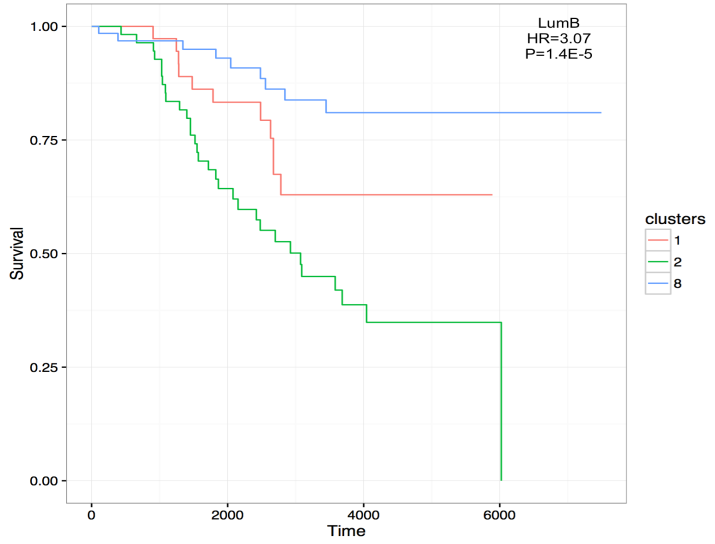

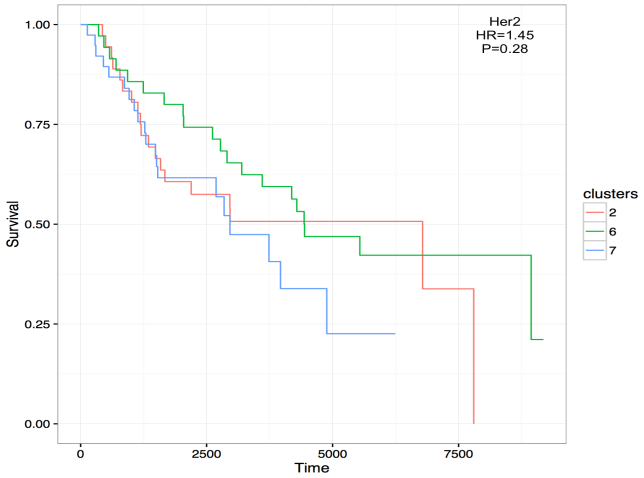

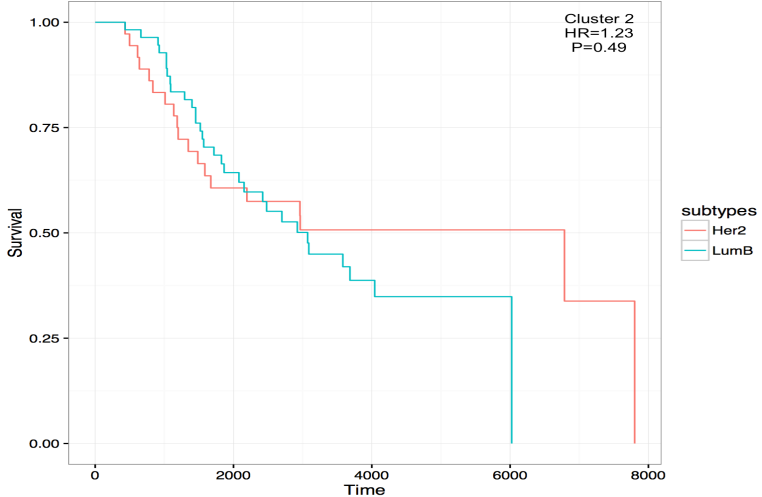

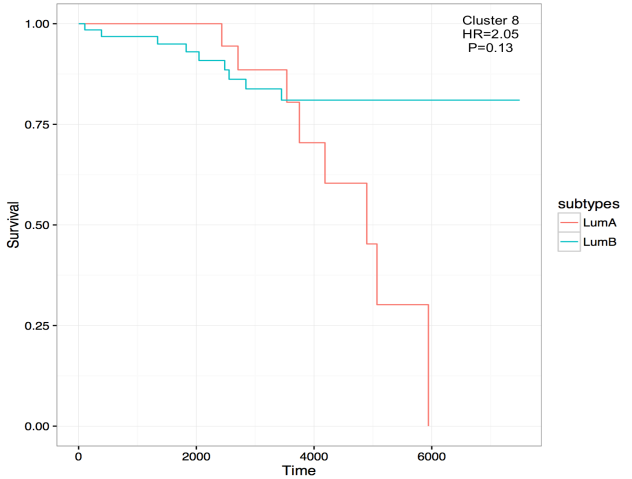

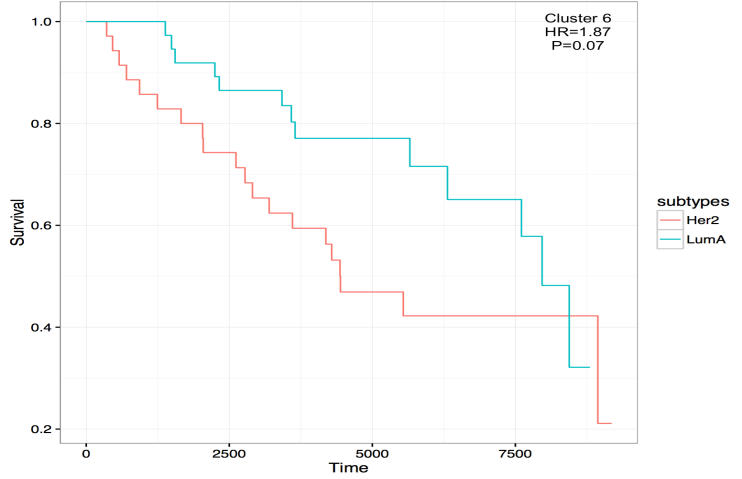

**Figure S9** - GI clustering accuracy measures relative to histopathological types.

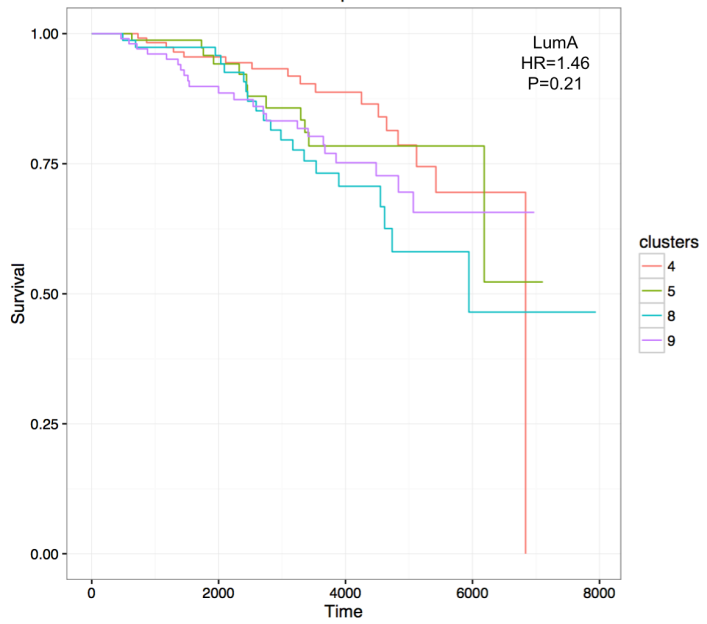

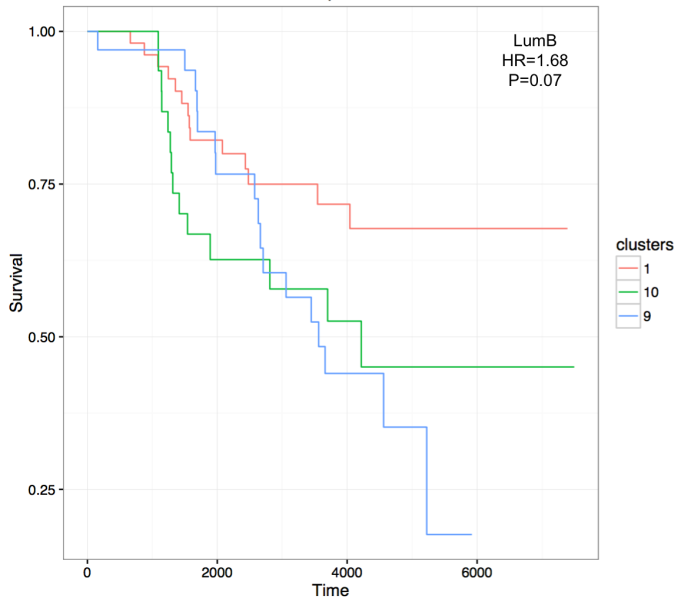

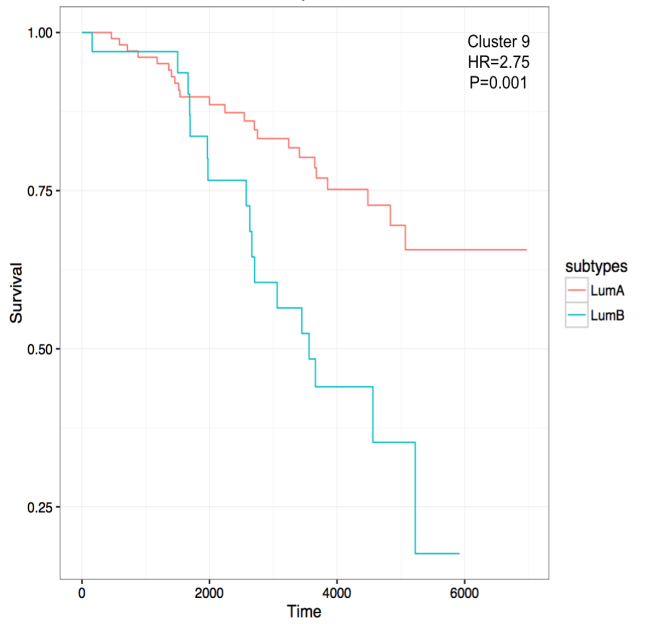

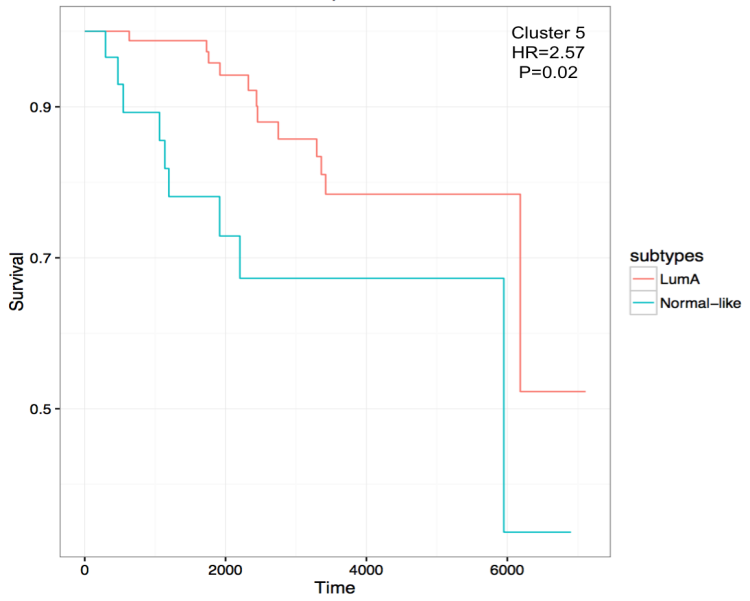

**Figure S10 -** METABRIC clustering accuracy relative to histopathological types

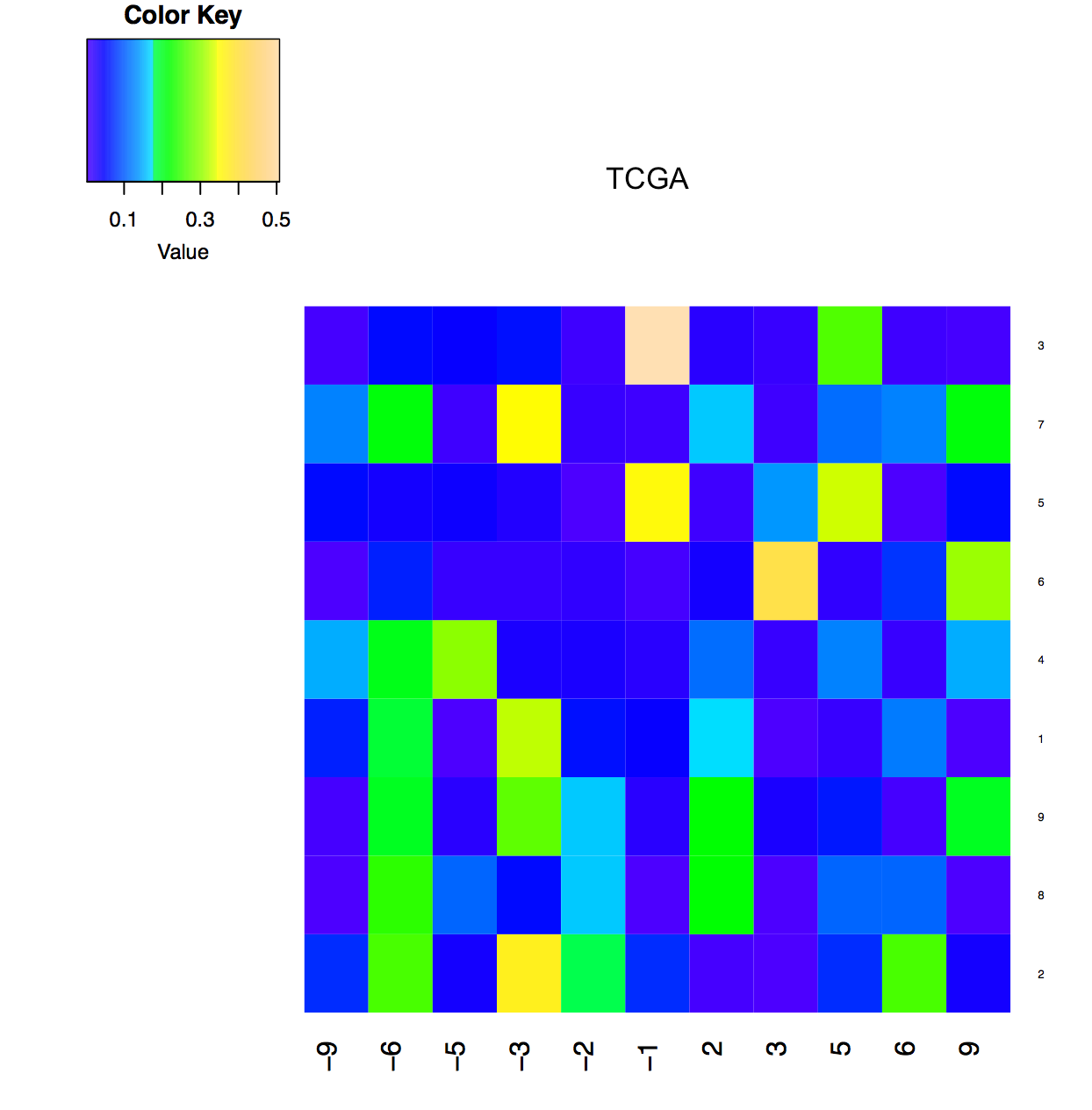

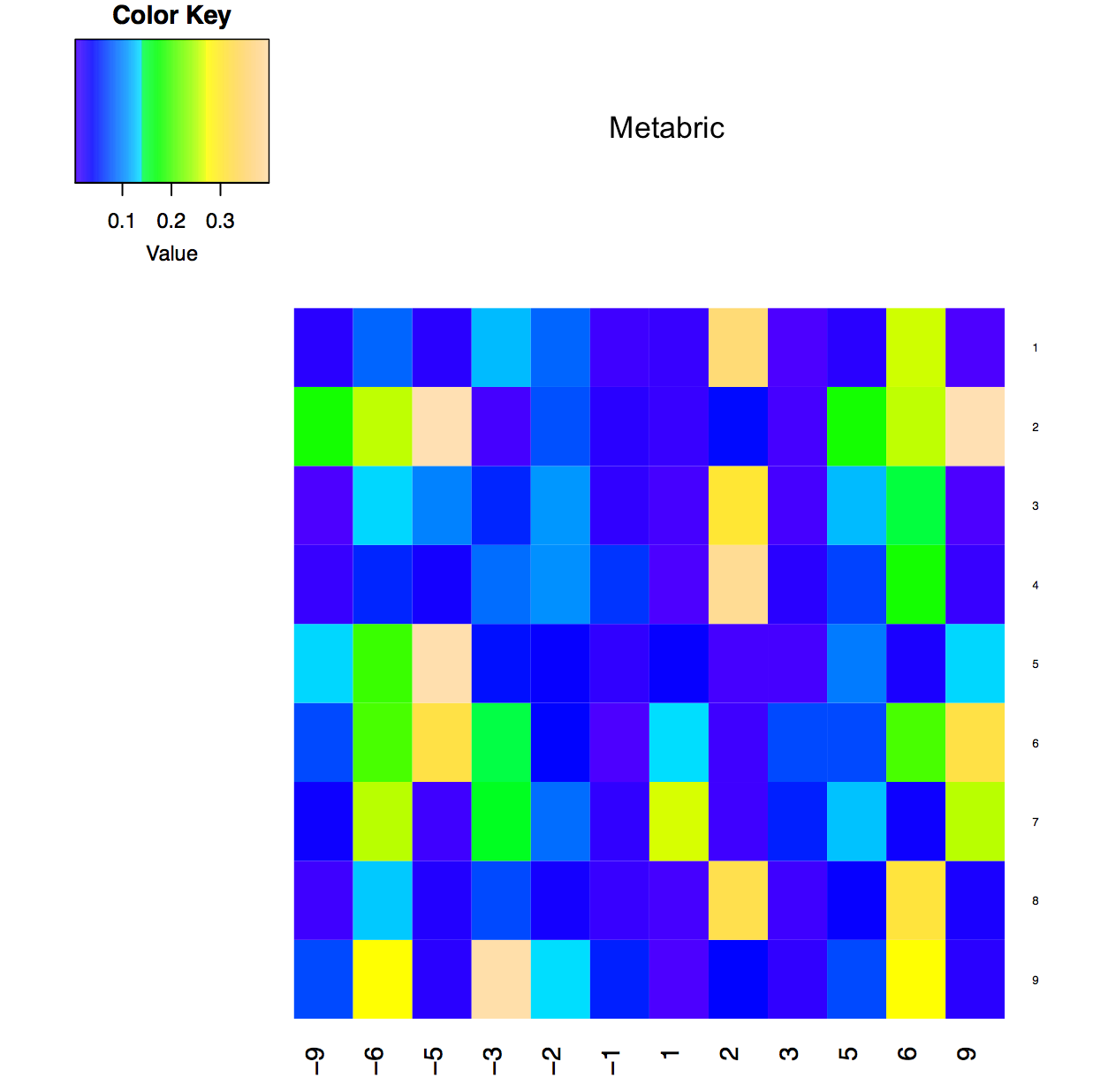

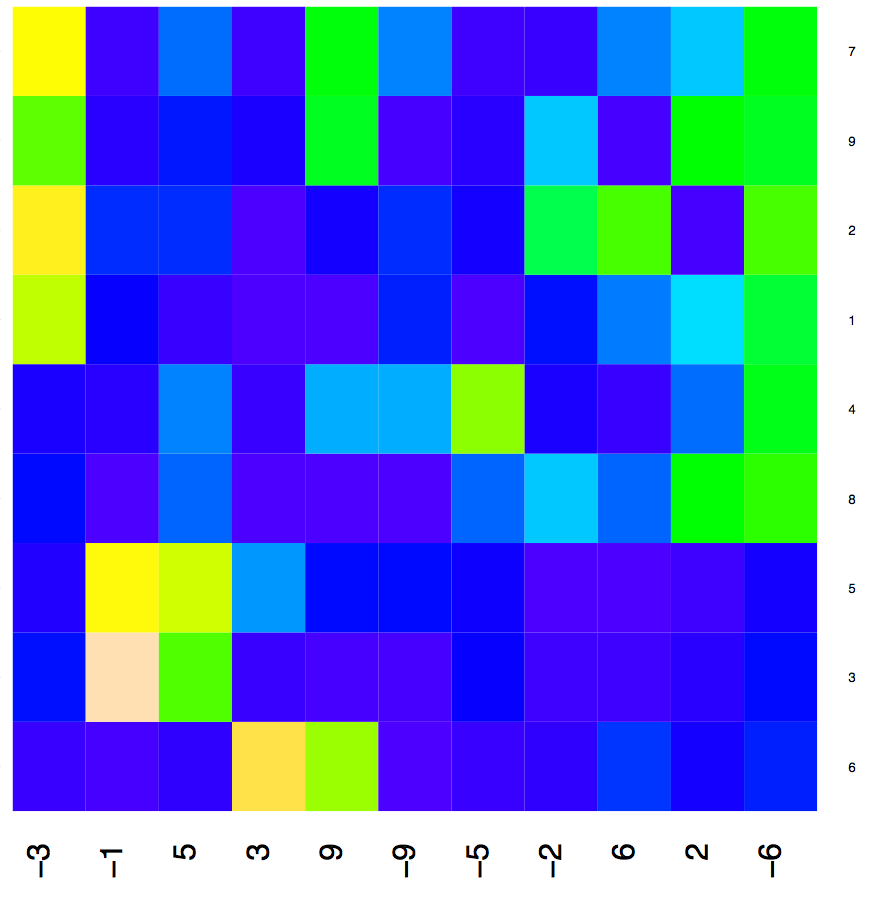

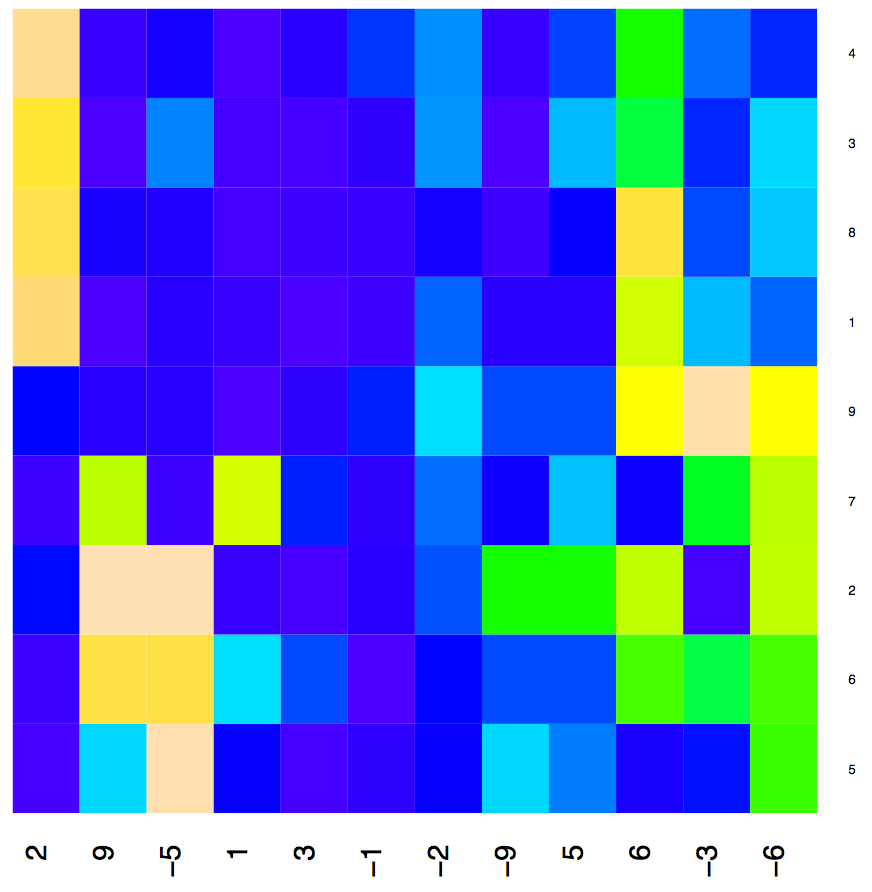

B

C

D

AC

B

**Figure S11** – GI-profiles of clusters revealed in the TCGA and the METABRIC breast cancer data and shows a high degree of consistency (Pearson correlation = 0.67, P-value = 1.1E-14). (A) METABRIC GI profiles (C) METABRIC GI profiles clustered by GI type. (B) TCGA GI profiles (D) TCGA GI profiles clustered by GI type.

**References**

Aran, D., Sirota, M., and Butte, A.J. (2015). Systematic pan-cancer analysis of tumour purity. Nat. Commun. *6*, 8971.

Bernard, P.S., Parker, J.S., Mullins, M., Cheung, M.C.U., Leung, S., Voduc, D., Vickery, T., Davies, S., Fauron, C., He, X., et al. (2009). Supervised risk predictor of breast cancer based on intrinsic subtypes. J. Clin. Oncol. *27*, 1160–1167.

Bilal, E., Dutkowski, J., Guinney, J., Jang, I.S., Logsdon, B.A., Pandey, G., Sauerwine, B.A., Shimoni, Y., Moen Vollan, H.K., Mecham, B.H., et al. (2013). Improving Breast Cancer Survival Analysis through Competition-Based Multidimensional Modeling. PLoS Comput. Biol. *9*.

Blazek, D., Kohoutek, J., Bartholomeeusen, K., Johansen, E., Hulinkova, P., Luo, Z., Cimermancic, P., Ule, J., and Peterlin, B.M. (2011). The Cyclin K/Cdk12 complex maintains genomic stability via regulation of expression of DNA damage response genes. Genes Dev. *25*, 2158–2172.

Chang, K., Creighton, C.J., Davis, C., Donehower, L., Drummond, J., Wheeler, D., Ally, A., Balasundaram, M., Birol, I., Butterfield, Y.S.N., et al. (2013). The Cancer Genome Atlas Pan-Cancer analysis project. Nat. Genet. *45*, 1113–1120.

Curtis, C., Shah, S.P., Chin, S.-F., Turashvili, G., Rueda, O.M., Dunning, M.J., Speed, D., Lynch, A.G., Samarajiwa, S., Yuan, Y., et al. (2012). The genomic and transcriptomic architecture of 2,000 breast tumours reveals novel subgroups. Nature *486*, 346–352.

Futreal, P.A., Coin, L., Marshall, M., Down, T., Hubbard, T., Wooster, R., Rahman, N., and Stratton, M.R. (2004). A census of human cancer genes. Nat. Rev. Cancer *4*, 177–183.

Gonzalez-Perez, A., Perez-Llamas, C., Deu-Pons, J., Tamborero, D., Schroeder, M.P., Jene-Sanz, A., Santos, A., and Lopez-Bigas, N. (2013). IntOGen-mutations identifies cancer drivers across tumor types. Nat. Methods *10*, 1081–1082.

Huang, X., Stern, D.F., and Zhao, H. (2016). Transcriptional Profiles from Paired Normal Samples Offer Complementary Information on Cancer Patient Survival – Evidence from TCGA Pan-Cancer Data. Sci. Rep. *6*, 20567.

Norton, N., Olson, R.M., Pegram, M., Tenner, K., Ballman, K. V, Clynes, R., Knutson, K.L., and Perez, E. a (2014). Association Studies of Fcγ Receptor Polymorphisms with Outcome in HER2+ Breast Cancer Patients Treated with Trastuzumab in NCCTG (Alliance) Trial N9831. Cancer Immunol. Res. *2*, 962–969.

Stephens, P.J., Tarpey, P.S., Davies, H., Van Loo, P., Greenman, C., Wedge, D.C., Nik-Zainal, S., Martin, S., Varela, I., Bignell, G.R., et al. (2012). The landscape of cancer genes and mutational processes in breast cancer. Nature *486*, 400–404.

Sun, Z., Shi, Y., Shen, Y., Cao, L., Zhang, W., and Guan, X. (2015). Analysis of different HER-2 mutations in breast cancer progression and drug resistance. J. Cell. Mol. Med. *19*, 2691–2701.

van de Vijver, M.J., He, Y.D., van’t Veer, L.J., Dai, H., Hart, A.A.M., Voskuil, D.W., Schreiber, G.J., Peterse, J.L., Roberts, C., Marton, M.J., et al. (2002). A gene-expression signature as a predictor of survival in breast cancer. N Engl J Med *347*, 1999–2009.
